# Cardiac neural crest lineage diversity and underlying gene regulatory networks revealed by multimodal analysis

**DOI:** 10.1101/2022.06.23.497419

**Authors:** Akiyasu Iwase, Yasunobu Uchijima, Daiki Seya, Mayuko Kida, Hiroki Higashiyama, Kazuhiro Matsui, Akashi Taguchi, Shogo Yamamoto, Shiro Fukuda, Seitaro Nomura, Takahide Kohro, Chisa Shukunami, Haruhiko Akiyama, Masahide Seki, Yutaka Suzuki, Youichiro Wada, Hiroyuki Aburatani, Yukiko Kurihara, Sachiko Miyagawa-Tomita, Hiroki Kurihara

## Abstract

Neural crest cells (NCCs), a multipotent stem cell population, contribute to cardiac development as a source of the outflow septum, vascular smooth muscle and semilunar valves. However, genetic programs underlying lineage diversification of cardiac NCCs remain largely unknown. Using single-cell (sc) and spatial transcriptomics, we demonstrate multiple NCC subpopulations with distinct gene expression signatures; smooth muscle(-like), non-muscle mesenchymal, and Schwann cell progenitor/melanoblast-like cells. Integrative scRNA-seq and scATAC-seq analyses predict lineage trajectories starting from immature NCCs, which bifurcate into smooth muscle(-like) and non-muscle mesenchymal lineages in association with hierarchical transcription factor networks. Combinatory analyses with Cre-mediated genetic lineage tracing characterize intermediate NCCs at the bifurcation as *Sox9*^+^/*Scx*^+^ tendon and cartilage progenitor-like cells with genetic programs, some of which are common to skeletal tissues whereas others are unique to cardiac NCCs. These findings provide a basis for understanding the roles of NCCs in cardiac development and pathogenesis particularly associated with calcification.

## Introduction

The neural crest (NC) is a migratory stem cell population that arises at the neural plate border in vertebrates and gives rise to various cell types including sensory and autonomic neurons, glial cells, adrenal chromaffin cells, pigment cells, and craniofacial skeletons^1–3^. In cardiac development, post-otic NCCs arising from rhombomeres 6–8 (known as ‘cardiac NCCs’) migrate into the 3rd, 4th, and 6th pharyngeal arches and the cardiac outflow tract, where they form the smooth muscle layer of the aorta and pulmonary artery, outflow septum, and the semilunar valves^4^. Removal of the cardiac NC in avian embryos results in cardiac malformations such as persistent truncus arteriosus^4–6^. In addition, NCCs derived from the preotic rhombomeres contribute to the coronary artery smooth muscle and a part of the semilunar valves, indicating that a wide range of the NC participates in cardiac development^7,8^. Recent innovations in single-cell analysis have facilitated studies on cell lineage trajectories and gene regulatory networks (GRNs). Soldatov et al. have demonstrated that cranial NCCs, unlike trunk NCCs, acquire ectomesenchyme potential before delamination from the neural tube depending on the transcription factor (TF) *Twist1*^9^. Similarly, the specification and differentiation of NCCs that contribute to cardiovascular formation are positionally determined before delamination, and also affected by environmental factors during migration^10^. Gandhi et al. identified *Tgif1* as a cardiac (postotic) NC-specific TF, whose deletion causes persistent truncus arteriosus in chick embryos^11^. Ectopic expression of *Tgif1*, together with other TFs *Ets1* and *Sox8*, can confer cardiovascular potential on trunk NCCs^11^. On the other hand, it remains largely unknown how sequential activation of GRNs enable diversification of NCC fates in the developing heart in response to spatiotemporal changes in environmental signals. Considering that the aorta, aortic valve, and coronary arteries are prone to calcification^12^, it is also important from a clinical viewpoint to explore the similarity and difference of the GRNs between cardiac (hereafter refers to both preotic and postotic) NCCs and cells of skeletogenic lineages including cranial NCC-derived ectomesenchymal cells.

To address these issues, we performed single-cell (sc)RNA-seq, spatial transcriptomic and multiomic analysis on embryonic cardiac NCCs from *Wnt1-Cre;R26-EYFP* mice, which are widely used to identify NCCs^13,14^. Together with immunostaining and lineage tracing using different Cre-expressing mouse lines, we identified diverse cell types derived from cardiac NCCs and predicted lineage trajectories. In particular, comparison of GRNs between cardiac NCCs and skeletogenic, especially teno-chondrogenic, progenitors provide an insight into the mechanisms how cardiac NCCs adopt differentiation programs into smooth muscle and mesenchymal cells such as valvular interstitial cells. The present findings may serve as a developmental basis for elucidating how pathological calcification occurs preferentially in the coronary artery, aortic wall, and aortic valve, to which cardiac NCCs highly contribute.

## Results

### Heterogeneity and lineage relationship of NCC populations in the mouse developing heart

To characterize the cardiac NCC lineages, we performed scRNA-seq on NCCs obtained from cardiac outflow tissues of *Wnt1-Cre;R26-EYFP* embryos using two different analytical platforms, the Fluidigm C1 and 10x Genomics Multiome systems (Fig. 1a). For scRNA-seq using the Fluidigm C1 system, EYFP^+^ NCCs were isolated from E11.5, E12.5, E14.5 and E17.5 samples by fluorescence-activated cell sorting (FACS) to yield a total of 309 cells (Supplementary Fig. 1a, b). After quality filtering, a dataset comprising 294 cells was subjected to unsupervised clustering analysis and projected onto the UMAP space, which separated these cells into two major compartments corresponding to early (E11.5, E12.5 and E14.5) and late (E17.5) developmental stages, and one minor population (Fig. 1b, Supplementary Fig. 1c-g). They were further separated into 13 clusters (fCs) with distinct transcriptomic profiles (Fig. 1c, Supplementary Fig. 1h, i, Supplementary Data 1). For 10x Genomics Multiome analysis, scRNA-seq and single-cell Assay of Transposase Accessible Chromatin sequencing (scATAC-seq) data were obtained from 236 and 198 NCCs at E14.5 and E17.5, respectively, after stringent quality control and normalization (Supplementary Fig. 2a-h). Unsupervised clustering with UMAP separated a total of 434 cells largely into two major and one minor compartments (Fig. 1d, Supplementary Fig. 2i). The two major compartments corresponded to smooth muscle-like and non-muscle mesenchymal phenotypes, which were grouped into 4 and 5 clusters (mCs), respectively (Fig. 1e-j, Supplementary Fig. 2j, k, Supplementary Data 2). Then, we combined both datasets and integrated them into a UMAP space based on canonical correlation analysis (CCA) dimensional reduction method after SCT normalization to analyze lineage diversity. The integrated data were divided into 8 clusters (intCs) (Fig. 1k, l, Supplementary Fig. 3a-c, Supplementary Data 3). Among Fluidigm C1 (fC) and 10x multiome (mC) clusters, fC2, fC4, mC6 and mC8 were split into separate groups in the integrated UMAP and were further subclustered for subsequent analysis (Fig. 1c, h, k, Supplementary Fig. 3e-h).

**Fig. 1.**
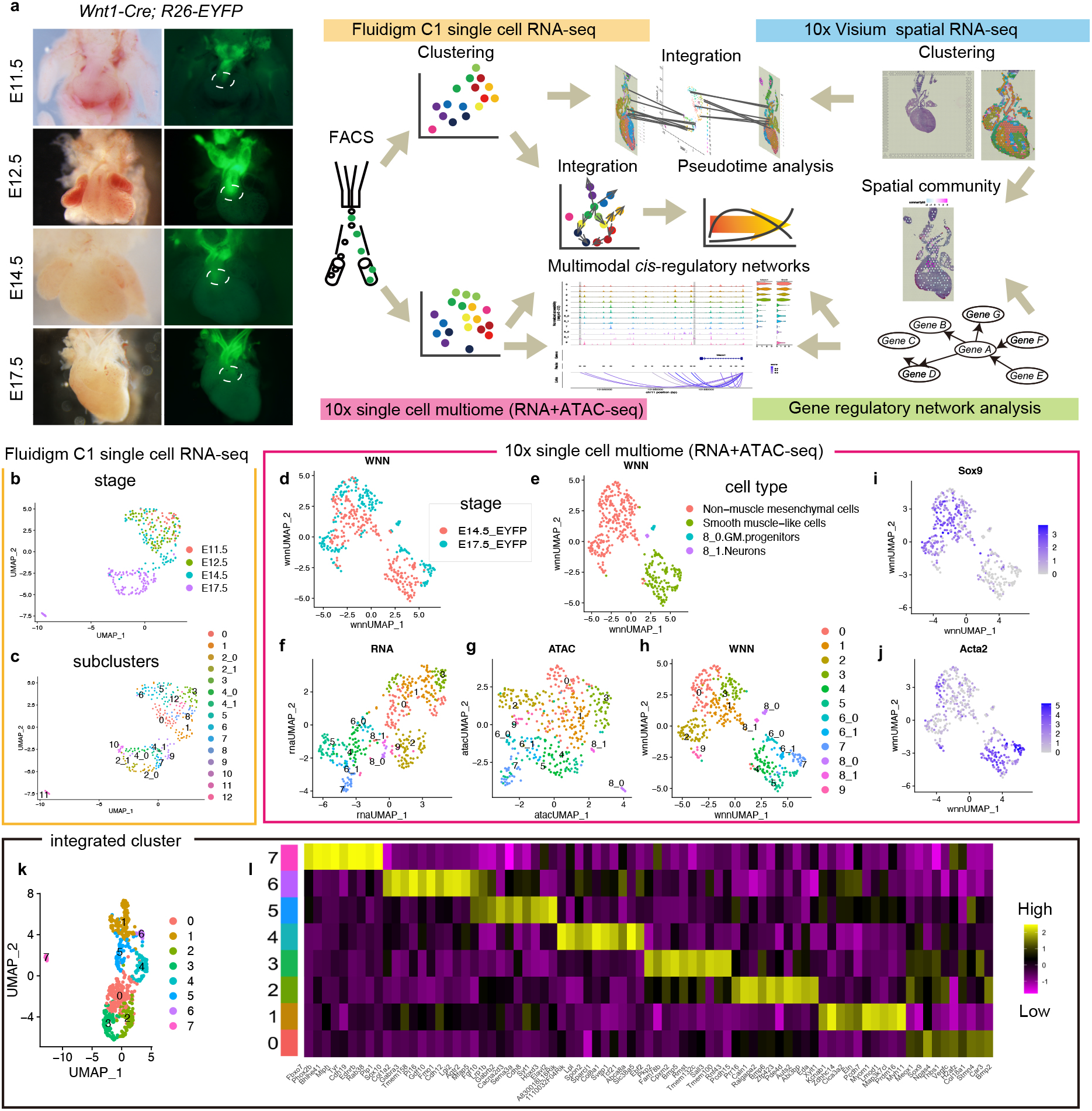
Heterogeneity of cardiac NCCs during heart development. (a) Workflow of single-cell and spatial genomic data analysis of cardiac NCCs. (b, c) UMAP plots of Fluidigm C1 scRNA-seq data, colored by embryonic stage (b) or cluster identity (c). (d-h) UMAP plots of scMultiome data, colored by embryonic stage (d), cell type (e) or cluster identity based on RNA-seq (f), ATAC-seq (g), or WNN (weighted-nearest neighbor) (h). (i-j) Feature plots of Sox9 (i) or Acta2 (j). (k) UMAP plots of integrated clusters by canonical correlation analysis of Fluidigm C1 and 10x Genomics scRNA-seq data. (l) Heatmap of differentially expressed genes (DEGs) among integrated clusters. DEGs were selected by the top 10 genes sorted by the value of average log2 fold change compared to the other cells. Expression values were averaged across integrated clusters.

IntC0 was largely composed of early-stage (especially E11.5 and E12.5) clusters (fC0, fC3, fC8, fC12, mC0 and mC1) (Fig. 1c, h, k, Supplementary Fig. 3g, h), representing an immature mesenchymal population characterized by high *Sox9* and *Meox1* expression (Fig. 1l, Supplementary Fig. 4a, b). Late-stage Fluidigm C1 and other 10x multiome non-muscle mesenchymal clusters were grouped into three subsets: intC2 (fC1 and fC9, *Bmp6*^high^), intC3 (fC4_1 and fC7, *Sall3*^high^) and intC4 (fC2_0, *Tcf21*^high^ and *Ebf2*^high^) (Fig. 1c, h, k, l, Supplementary Fig. 3g, h, 4c-f). IntC4 also included mC9 characterized by pericyte marker *Abcc9* (Fig, 1h, k, Supplementary Fig. 2k, 4g). Clusters with smooth muscle-like characteristics (e.g. fC4_0, fC10 and many of smooth muscle-like mC components) constituted intC1 with highly expressed mature smooth muscle markers (*Myh11, Lmod1* and *Myom1*) (Fig. 1c, h, k, Supplementary Fig. 3g, h, 4i-k). IntC1 contained two subtypes as represented by mC7_0 (*Myh11*^+,^ *Kcnj8*^-^) and mC6_1 (*Myh11*^+^, *Kcnj8*^+^) (Supplementary Fig. 2l, 4i, l), which may correspond to smooth muscle cells (SMCs) of the great vessel and coronary artery SMCs, respectively, as indicated by Chen and colleagues (see Discussion)^15^. Remarkably, intC0 and intC1 were connected by intC5 characterized by *Sema3a* expression, which is mainly composed of fC5 and fC6 (Fig. 1c, k, l, Supplementary Fig. 4m). IntC6, mainly composed of fC2_1 and mC6_0, was continuous with intC1 and intC5 and characterized by *Dlk1*, *Cxcl12* and *Mfap5* expression (Fig. 1c, h, k, l, Supplementary Fig. 1i, 2k, 3g, h, 4n-p). Apart from these clusters, intC7 represented a minor population composed of fC11 and mC8_0 and was characterized by markers of Schwann cell precursors (e.g. *Sox10* and *Plp1*) or melanocytes (e.g. *Tyr*) (Fig. 1c, h, k, l, Supplementary Fig. 1i, 2k, 3g, h, 4q-s).

Then we performed pseudotime analysis using Monocle3 to infer lineage trajectories. IntC7 was excluded from this analysis because it was categorized into a different partition (Supplementary Fig. 3d). The root node of the trajectories was determined by using RNA velocity analysis of the early-stage Fluidigm C1 dataset, which showed two general flows starting from fC3, mainly composed of E11.5 NCCs characterized by early endocardial cushion mesenchymal markers (e.g. *Twist1*, *Sox9*, *Vcan* and *Hapln1*), towards fC6 and fC1 for each (Fig. 2a, Supplementary Fig. 5a-d), suggesting a bifurcation from immature cushion-forming NCCs toward *Acta2*^high^ smooth muscle-like and *Acta2*^low/–^ non-muscle mesenchymal differentiation (Fig. 2a, Supplementary Fig. 5e). Accordingly, we obtained presumptive pseudotime trajectory branching from the root node within intC0 containing fC3 into five different destinations, corresponding to intC1, intC2, intC3, intC4 and intC6) (Fig. 2b, c, Supplementary Data 3).

**Fig. 2.**
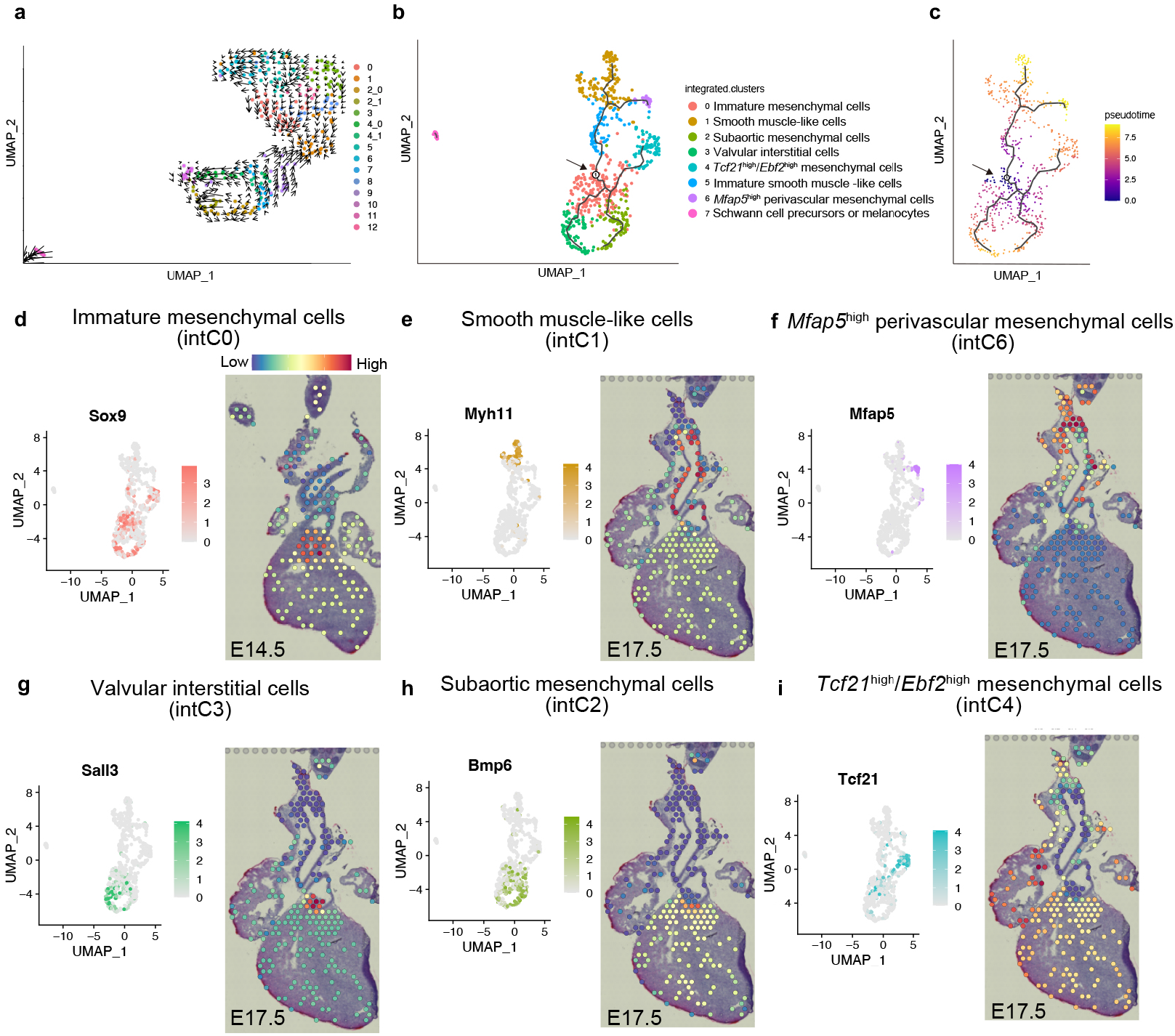
Lineage trajectories of cardiac NCCs with spatial identity. (a) RNA velocity flow embedded on the UMAP of Fluidigm C1 data. (b, c) Pseudotime trajectories (black lines) from the root node (arrows) embedded on the UMAP of integrated clusters, colored by cluster identity (b) or pseudotime (c). (d-i) Projection of integrated clusters (intCs) onto the Visium data (right) with marker gene expression plots on the UMAP (left) to predict their spatial identities. The color bar above each Visium image indicates the probability of the spatial prediction.

### Spatial allocation of NCC clusters by integration of single-cell and spatial transcriptomic analysis

To obtain information on the spatial distribution and connectivity of each cluster revealed by scRNA-seq, we performed spatial transcriptomic analysis on sections of E14.5 and E17.5 *Wnt1-Cre;R26-EYFP* mouse hearts and surrounding tissues using the 10x Genomics Visium platform (Supplementary Fig. 6a-l). Clustering analysis classified datasets of 702 (E14.5) and 1428 (E17.5) spots into 23 and 32 clusters, respectively, which were designated as early and late Visium clusters (evCs and lvCs) (Supplementary Fig. 6b, c, h, i, Supplementary Data 4). After NCC-containing *EYFP*-positive spots were extracted from the sequence reads (Supplementary Fig. 6e, k), each single-cell cluster (intC) was allocated to these spots with probability estimated from single-cell and spatial transcriptomic profiling data (Fig. 2d-i, Supplementary Fig. 7a-d, 8a-d).

When intCs were projected onto E14.5 Visium datasets, intC0 and intC3 were allocated to the semilunar valves and endocardial cushion region (evC4) (Fig. 2d, g, Supplementary Fig. 6b, 7a, b), whereas intC1 was allocated to the great vessel walls and surrounding tissues (evC3) (Fig. 2d, g, Supplementary Fig. 6b, 7a, b). Compared to intC0, intC3 signals were preferentially found in the semilunar valves (Fig. 2d, g, Supplementary Fig. 7d), which was more obvious in the projection onto E17.5 Visium datasets (lvC31) (Supplementary Fig. 6h). IntC1 was also allocated to the aorta and pulmonary artery at E17.5 (late Visium cluster (lvC)5 and lvC12) (Fig. 2e, Supplementary Fig. 6h, 7c, d). IntC4 and intC6 were allocated to tissues surrounding the great vessels (lvC8, 9 and 16), with intC6 closer to the intC1^+^ spots in the great vessel wall (Fig. 2f, i, Supplementary Fig. 6h, 7c, d). Although intC5 was not allocated to any specific *EYFP*-positive areas (Supplementary Fig. 7a-d), fC6, a major constituent of intC5, was allocated to the great vessel walls and surrounding tissues (evC3) at E14.5 (Supplementary Fig. 3g, 8a, b), consistent with the idea that intC5 represents an intermediate stage of smooth muscle(-like) differentiation. Similarly, intC2 did not correspond to any specific *EYFP*-positive areas (Supplementary Fig. 7a-d), but its major constituents fC1 and fC9 were allocated to the endocardial cushion region (evC4 and lvC18), corresponding to the subaortic mesenchymal condensation enriched in NCCs (Fig. 2h, Supplementary Fig. 3g, 6h, 8a-d). Taken together, the five distinct NCC lineages inferred from pseudotime analysis of the integrated data are likely to mainly represent smooth muscle(-like) (intC1), subaortic mesenchymal (intC2), valvular interstitial (intC3), *Tcf21*^high^/*Ebf2*^high^ mesenchymal (intC4) and *Mfap5*^high^ perivascular mesenchymal (intC6) lineages (Fig. 2d-i). It is worth mentioning that *Tcf21* was also expressed in some cells belonging to intC0 and intC3, but *Ebf2* was not (See Discussion) (Supplementary Fig. 4e, f).

### Inference of GRNs in NCC differentiation

To infer GRNs underlying cardiac NCC differentiation, we performed nonparametric Bayesian network analysis on the Fluidigm C1 scRNA-seq dataset except for fC11 by using the NNSR algorithm^16^, taking advantage of deeper sequencing than the 10x multiome dataset (Fig. 3a, Supplementary Fig. 9a). Based on 269,646 potential regulatory links between TFs and target genes extracted from SCENIC^17^, we identified 18,655 links with estimated frequencies higher than the cutoff threshold. From these links, genes annotated as ‘transcription factor’ in SCENIC were extracted, resulting in a network composed of 560 gene nodes and 1588 edges (Supplementary Fig. 9b). The yielded network was divided into 109 communities with some overlapping by the Linkcomm algorithm^18^ (Supplementary Fig. 9b). We then constructed subnetworks in which TF(s) belonging to any of the communities are connected to nodes of the first edge, referred to as “overall communities (OCs)”, and estimated their intensity per cell (scRNA-seq) or spot (Visium) (Fig. 3a, b, Supplementary Fig. 11, Supplementary Data 5).

**Fig. 3.**
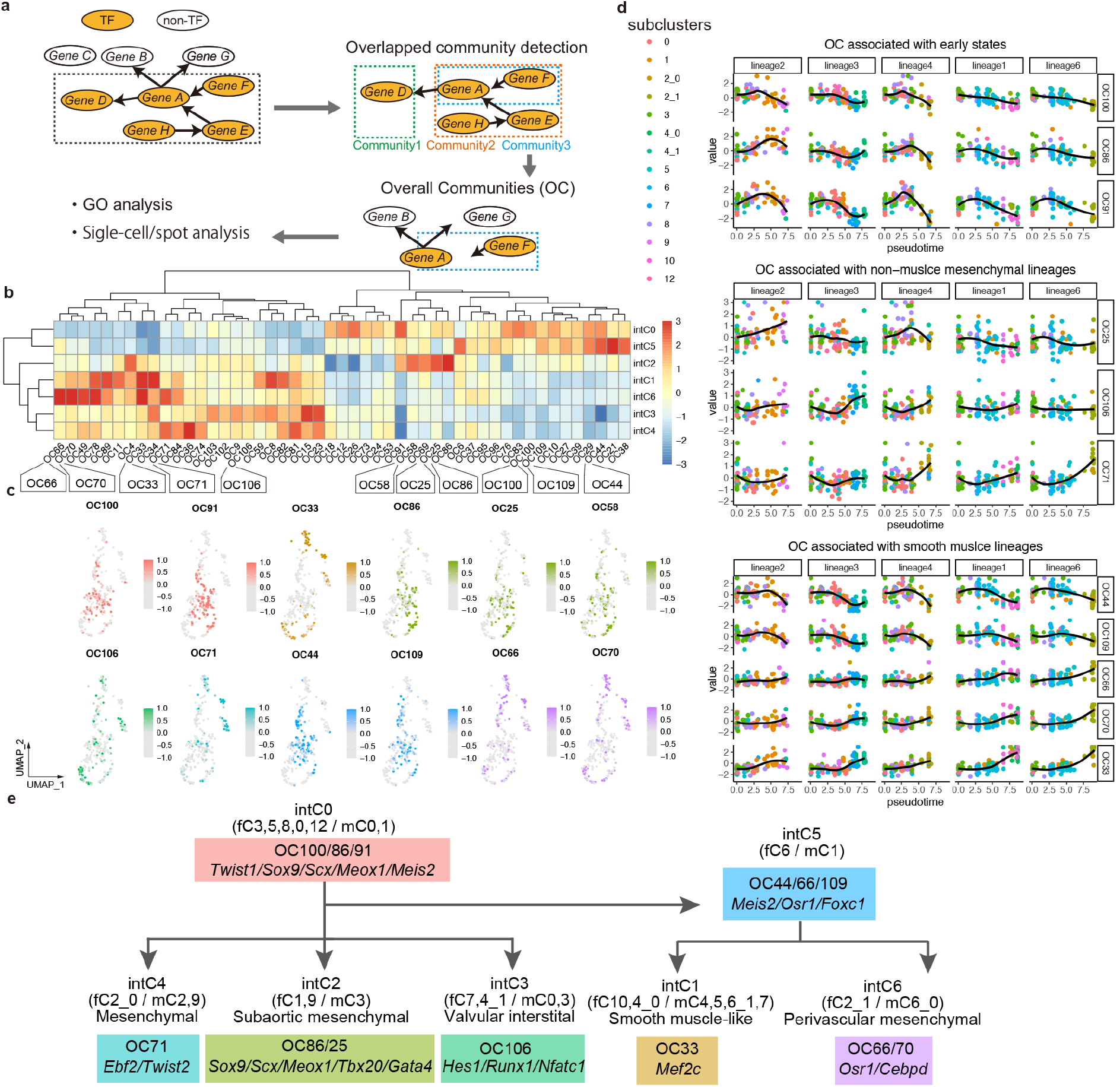
GRNs involved in cardiac NCC fate determination. (a) Scheme of GRN analysis based on Fluidigm C1 scRNA-seq data. Subnetwork of TFs surrounded by dot line was divided into communities with some overlapping. TF(s) belonging to any of the communities are connected to nodes of the first edge, referred to as “overall communities (OCs)”. (b) Heatmap showing the expression of OCs across the integrated clusters. The color indicates z-scored value across OCs. (c) Representative OC expression on the UMAP plots, colored by cluster identity corresponding to (e) with intensity of expression levels. (d) Distribution of OC along the pseudotime shown in Figure 2c. Changes in expression levels of OCs associated with early states (intC0) (top), non-muscle mesenchymal lineages (intC2, intC3 and intC4) (middle) and smooth muscle(-like) lineages (intC1 and intC6) (bottom) are shown. Horizontal axes indicate pseudotime set as Fig. 2c. Lineage2, intC0 to intC2; lineage3, intC0 to intC3; lineage4, intC0 to intC4; lineage1, intC0 to intC1 through intC5; lineage6, intC0 to intC6 through intC5. Each color indicates cluster identity corresponding to (e). (e) Prediction of integrated lineages with OCs and key TFs.

The earliest cardiac NCC population around the root node within intC0 were characterized by GRNs involving TFs implicated in NC-derived ectomesenchyme and/or endocardial cushion development, such as OC 100 (containing *Twist1*, *Prrx2*, *Sox4* and *Sox11*) and OC91 (*Foxp2*, *Meis2*, *Mecom*, *Id2*, *Pbx3*, *Sox4* and *Sox5*) (Fig. 3b, c, Supplementary Fig. 9c, e), consistent with gene ontology (GO) terms like ‘outflow tract morphogenesis’ (Supplementary Fig. 10). As development progressed, OC86 (*Sox9*, *Scx*, *Meox1, Meis2, Tbx20*, *Foxp2* and *Sox4*) became prevalent especially in intC2 (Fig. 3b, c, Supplementary Fig. 9d). whose spatial identity was confirmed by allocation to the outflow cushion mesenchyme (evC4) in the Visium clustering (Supplementary Fig. 5b, 11a-c). IntC2 was further characterized by upregulation of OC25 (*Tbx20* and *Gata4*) (Fig. 3b, c, Supplementary Fig. 9f), which was mapped to the Visium plots in the subaortic region (Supplementary Fig. 11d). Other non-muscle mesenchymal lineages intC3 and intC4 were characterized by OC106 (*Hes1*, *Runx1* and *Nfatc1*) and OC71 (*Ebf2, Klf4, Plagl1* and *Twist2*), respectively (Fig. 3b, c, Supplementary Fig. 9g, h, 11e,f).

In smooth muscle-directed lineages, the intermediate cluster intC5 shared OC44 (*Pbx2*, *Meis2* and *Smarca4*) with intC0, indicating a putative smooth muscle-directed GRN (Fig. 3b, c, Supplementary Fig. 9i), consistent with GO terms of smooth muscle development and coronary vasculature development (Supplementary Fig. 10). OC44 was enriched in the great artery and cushion mesenchyme region in the Visium data (Supplementary Fig. 11g), the latter of which might represent coronary arteries. IntC1 and intC6, OC66 (*Osr1*, *Osr2*, *Hlx* and *Klf15*), OC70 (*Osr1* and *Cebpd*) and OC 109 (*Foxc1*) were enriched (Fig. 3b, c, Supplementary Fig. 9j-l, 11h-j). Because Osr1 has been reported to suppress *Sox9* expression^19^, upregulation of Osr1 may facilitate smooth muscle(-like) differentiation by suppressing Sox9-dependent GRN. IntC1 and intC6 were also characterized by OC33 (*Mef2c, Cepbd, Fos*, *Klf2* and *Egr1*) (Fig. 3b, c, Supplementary Fig. 9m), which was associated with GO terms of muscle structure development and negative regulation of cell proliferation and was enriched in the great artery of smooth muscle and perivascular mesenchymal regions (Supplementary Fig. 10, 11k). Notably, *Cebpd* connected to *Mfap5*, serving as a marker for intC6, in OC70, which was annotated with GO terms involved in extracellular matrix organization and fibroblast migration (Fig. 3b, c, Supplementary Fig. 9k, 10), indicating intermediate characteristics between smooth muscle cells and fibroblasts.

Dynamics of GRNs represented by changes in OC levels along the pseudotime from immature mesenchyme to each of the five lineages and schematic summary are given in Fig. 3d and 3e with Supplementary Data. 3.

### Inference of multimodal *cis*-regulatory networks in NCC differentiation

To validate GRNs inferred from the Fluidigm C1 scRNA-seq dataset, we performed multimodal *cis*-regulatory network analysis based on the epigenomic and transcriptomic data from the 10x Genomics scMultiome platform (RNA-seq and ATAC-seq). First, we estimated chromatin accessibility of TF-binding motifs by chromVAR^20^ and extracted TFs involved in different processes of NCC differentiation. Average accessibility of TF-binding motifs was variable among integrated clusters and was largely correlated with the expression level for many TFs in OCs characterizing cardiac NCC lineages (Fig. 4a, Supplementary Fig. 12, Supplementary Data. 2). Non-muscle mesenchymal clusters (IntC0, 2, 3 and 4) were enriched for accessible binding motifs of TFs Sox9, Meox1, Tbx20, Lef1, Gata4, Nfatc1 and Runx1 (Fig. 4a, h, i, Supplementary Fig. 12a-f), whereas smooth muscle(-like) clusters (intC5, 1 and 6) showed high accessibility of Foxc1, Mef2c and Tead3-binding motifs (Fig. 4a, n, o, Supplementary Fig. 12k, l). In addition, IntC4 was enriched for accessible Ebf2, Tcf21, and Ets1-binding motifs (Fig. 4a, Supplementary Fig. 12g-j). Interestingly, two different Meis2-binding motifs MA07774.1 and MA1640.1 showed opposing accessibility in mesenchymal and smooth muscle(-like) clusters, respectively (Fig. 4a, c-e). Given that Meis2, a component common to OC86/91 and OC44, which were GRNs characterizing non-muscle mesenchymal and smooth muscle(-like) lineages, respectively (Fig. 4a, c-e, Supplementary Fig. 4u, 9d, e, i), Meis2 may be involved in distinct lineages with different binding motifs.

**Fig. 4.**
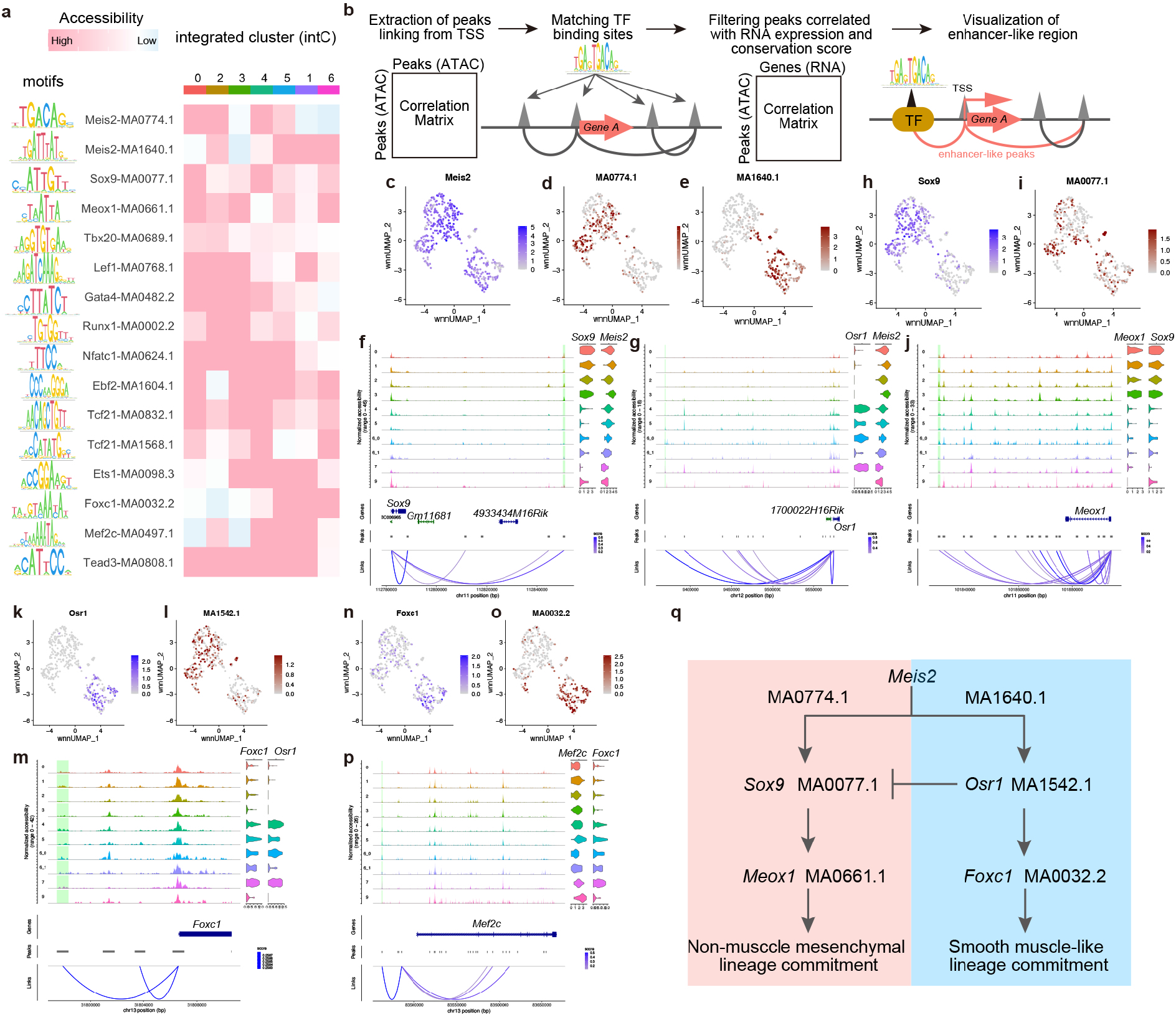
*Cis*-regulatory networks in cardiac NCC development. (a) Heatmap of motif activities. Motif activities were averaged across integrated clusters. Value of accessibility was averaged across integrated clusters. (b) Workflow for identifying *cis*-regulatory networks by scMultiome data including scATAC-seq and scRNA-seq. (c-p) RNA expression (c, h, k, n) and TF motif accessibility (d, e, i, l, o) on the UMAP plots with coverage plots (left) from scATAC-seq data and RNA expression of violin plots (right) from scRNA-seq data (f, g, j, m, p). For coverage and violin plots, each row indicates mC, with the enhancer-like region highlighted with green. (q) Scheme of *cis*-regulatory networks during lineage bifurcation in cardiac NCC development.

To further explore regulatory hierarchy of TFs, multimodal *cis*-regulatory networks were constructed using the 10x Multiome dataset by presuming binding motifs corresponding to enhancer-like regions that are positively correlated with gene expression among co-accessible peaks associated with transcription start sites (TSSs) and are highly conserved among Euarchontoglires (Fig. 4b, Supplementary Data 6). Notably, the MA0774.1 Meis2-binding motif was found in the mesenchymal open chromatin region correlated to the TSS of *Sox9* (Fig. 4f). *Sox5, Tbx20,* and *Runx1*, components of mesenchymal communities such as OC25, 86, 91, and 106 (Supplementary Fig. 8d-g), were also targeted by MA0774.1 (Supplementary Fig. 13a-d). In mesenchymal clusters, Sox9-binding motifs were found in the open chromatin regions correlated to the TSS of *Meox1* (Fig. 4h-j), consistent with the directional connections among *Meis2*, *Sox9* and *Meox1* in OC86 (Supplementary Fig. 9d).

Another Meis2 motif MA1640.1 was found in open chromatin regions related to the TSS of *Osr1*, whose expression was enriched in smooth muscle(-like) clusters (Fig. 4a, c, e, Supplementary Fig. 4v). Osr1 was predicted to bind to the enhancer-like region of *Foxc1*, whose expression and binding motif accessibility were enriched in mC4 and mC5 largely belonging to intC1 (Fig. 4a, k-m, Supplementary Fig. 4w). Foxc1 has been reported to be required for smooth muscle differentiation in zebrafish^21^, suggesting the Osr1-Foxc1 axis may be involved in smooth muscle differentiation. In addition, Foxc1-binding motifs were found in *Osr1*, *Mef2c*, and *Tead3*, which are selective for smooth muscle-like lineages (Fig. 4a, n-p, Supplementary Fig. 13e, f), indicating a positive feedback loop between Foxc1 and Osr1. Moreover, it is known that Mef2 and Tead function upstream of *Myocd*, a key regulator of smooth muscle genes, in a cooperative manner^22^. *Mef2c* belongs to OC33 (Supplementary Fig. 9m), suggesting a commitment to smooth muscle differentiation, consistent with the results of GRN analysis of Fluidigm C1 scRNA-seq data (Fig. 3b-e). In summary, the analysis of GRNs and multimodal *cis*-regulatory networks revealed a mode of differentiation from non-muscle mesenchymal state to smooth muscle(-like) state with changes in chromatin accessibility involving different Meis2-bindig motifs (Fig. 4q).

### Involvement of Sox9 and Scx in cardiac NCC lineages

Among TFs involved in GRNs acting in cardiac NCC differentiation, *Sox9* and *Scx* in OC86 were expressed at early stages around bifurcation into smooth muscle and non-muscle mesenchyme lineages (Supplementary Fig. 4a, t, 5a, f, 11b). We then examined the expression patterns of Sox9 and Scx by using *Sox9-EGFP* and *Scx-Tomato* mice (Fig. 5a-i). At E12.5, Sox9 expression represented by EGFP signals were found in the endocardial cushion and its derivatives including semilunar valve primordia, which partially overlapped with SM22α and αSMA expression (Fig. 5b, c). Within these Sox9-positive regions, Scx expression represented by tdTomato was found in and around mesenchymal condensation (Fig. 5c). Scx expression was also found in great vessel walls and aorticopulmonary septal mesenchyme partially overlapping with smooth muscle markers (Fig. 5b). At E14.5 and E17.5, Sox9 expression was sustained in endocardial cushion derivatives with high expression of Scx in the mesenchymal condensation and aortic and pulmonary roots, especially in the fibrous interleaflet triangles (Fig. 5e, f, h). By contrast, Scx expression was relatively low in the semilunar valve leaflets at E17.5 (Fig. 5h). The Myh11-labeled smooth muscle layer of coronary arteries, which is also contributed to by NCCs, was negative for Sox9 and Scx expression (Fig. 5i). In the mesenchymal condensation mainly composed of NCCs, type 2 collagen (Col2a1)^23^, known as cartilage-specific collagen downstream of Sox9, was also highly expressed, as evidenced by immunostaining and scRNA-seq data (Fig. 5j, Supplementary Fig. 4x). These findings suggest that chondrocyte-like genetic program operates in the cushion mesenchymal NCCs.

**Fig. 5.**
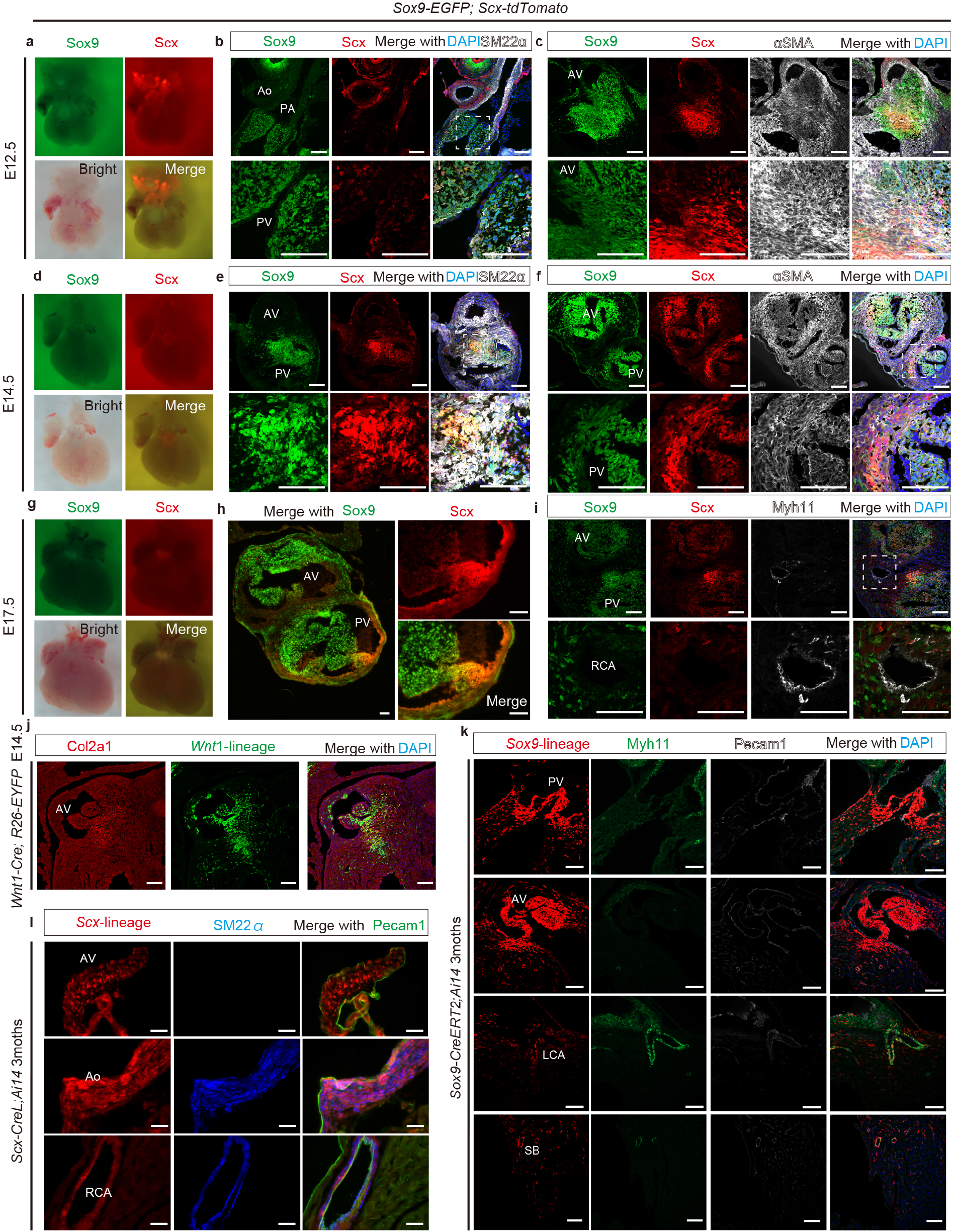
*Sox9^+^*/*Scx^+^* cells and their descendants in the developing heart. (a-i) Expression patterns of reporter genes in *Sox9-EGFP;Scx-tdTomato* mice. Whole-mount images (a, d, g) and immunostaining for Sox9-EGFP and Scx-tdTomato (b, c, e, f, h, i) with co-staining for SM22α (b, e), αSMA (c, f) or Myh11 (i) at E12.5 (a-c), E14.5 (d-f) and E17.5 (g-i). Boxed areas are magnified in lower panels. (j) Immunohistochemistry of Col2a1 and EYFP (*Wnt1*-lineage) in *Wnt1-Cre;R26-EYFP* mice at E14.5. (k) Immunohistochemistry of Myh11, Pecam1 and tdTomato (*Sox9*-lineage) in E17.5 *Sox9-CreERT2;Ai14* mice treated with tamoxifen at E11.5. (l) Immunohistochemistry of SM22α, Pecam1 and tdTomato (*Scx*-lineage) in *Scx-CreL;Ai14* mice at 3 months. Ao, aorta; AV, aortic valve; LCA, left coronary artery; PA, pulmonary artery; PV, pulmonary valve; RCA, right coronary artery; SB septal branch of coronary artery. Nuclei are stained with DAPI. Scale bars, 50 μm.

Then we tested whether Scx-negative valve leaflets and coronary artery smooth muscle differentiate through a *Sox9*^+^/*Scx*^+^ intermediate state by using Cre reporter mice (Fig. 5k, l). In *Sox9-CreERT2;Ai14* mice treated with tamoxifen at E11.5, Cre-mediated recombination was broadly detected in the wall of aortic and pulmonary roots, semilunar valve leaflets, the smooth muscle layers of coronary arteries and surrounding mesenchymal cells at E17.5 (Fig. 5k). *Scx-CreL; Ai14* mice also labeled the aortic wall, semilunar valve leaflets and coronary artery SMCs (Fig. 5l). These results indicate that the most of cardiac NCC-derivatives differentiate through a *Sox9*^+^/*Scx*^+^ intermediate state.

### Characterization of *Sox9*^+^/*Scx*^+^ intermediate cell population

To characterize *Sox9*^+^/*Scx*^+^ cells in the cardiac NCC lineage, we extracted high expressors of both genes from the Fluidigm C1 scRNA-seq dataset containing early-stage cardiac NCCs (Supplementary Fig. 14a, d, e). These *Sox9*^high^/*Scx*^high^ cells were divided into 4 subclusters (sC0 to sC3) largely according to their parent fC clusters (Fig. 6a, Supplementary Fig. 14b, c). The earliest subcluster sC3 differentially expressed *Twist1*, *Vcan* and *Hapln1*, as with fC3 (Supplementary Fig. 5b-d, 14f-h, Supplementary Data. 7). sC2 and sC1 corresponded to non-muscle mesenchymal (fC1/fC9) and smooth muscle(-like) (fC5/fC6) lineages, respectively (Supplementary Fig. 14c), which are considered to stem from sC3 (Fig. 2a). Notably, *Sox9*- and *Scx*-containing OC86 was enriched in sC2 and, to a lesser extent, sC3, indicating that GRNs involving Sox9 and Scx is active in the mesenchymal lineage (Fig. 6b, Supplementary Fig. 14i). *Sox9-* and *Scx*-expressing cells were also found in late-stage subclusters of both lineages (sC0) (Supplementary Fig. 14c).

**Fig. 6.**
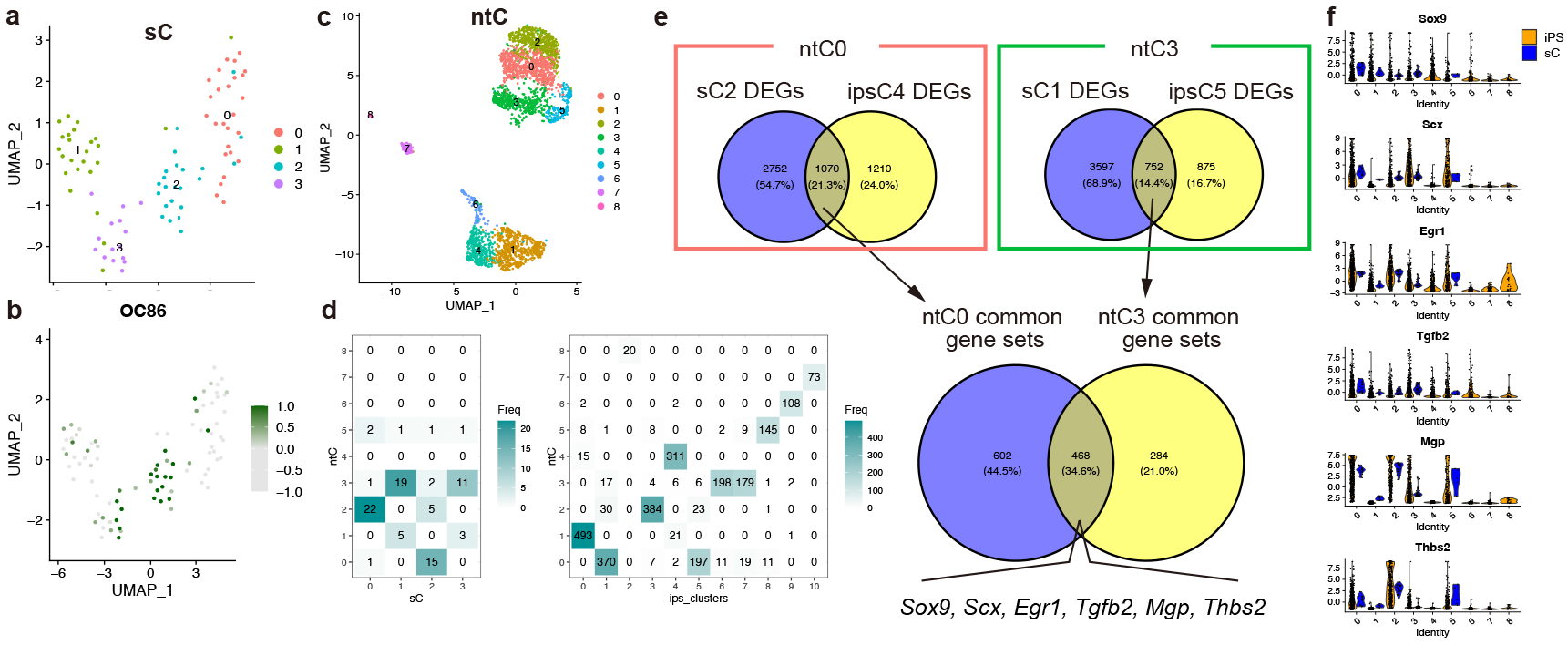
*Sox9*^+^/*Scx*^+^ cardiac NCCs and tenocyte progenitors share common gene expression signatures. (a, b) UMAP plots of *Sox9*^high^/*Scx*^high^ NCC subclusters, labeled by subcluster identity (sC) (a) and OC86 levels (b). (c) UMAP plots of combined scRNA-seq data from fC clusters and iPS-derivatives^25^, colored by cluster identity (ntC). (d) Cross tables between ntC and sC (left) or iPS clusters in the original report^25^ (right). Color bars indicate the frequency. (e) Venn diagrams comparing DEGs between sC and iPS data sets in ntC0 (left), ntC3 (right), and common gene sets (bottom). (f) Violin plots of expression profiles of common gene sets in the integrated sC and iPS data.

In musculoskeletal development, *Sox9*^+^/*Scx*^+^ cells serve as progenitors for tenocytes, ligamentocytes and chondrocytes^24^. To compare the nature of *Sox9*^+^/*Scx*^+^ cells between two different lineages, we integrated the Fluidigm C1 scRNA-seq dataset with a published scRNA-seq dataset for tenocyte differentiation from induced pluripotent stem cells (iPSCs)^25^ (Supplementary Fig. 14j-o, Supplementary Data. 7). As a result, sC1/sC3, sC2, and sC0 were co-clustered with iPSC-derived cells at different differentiation stages (NCC-tenocyte cluster (ntC) 3, ntC0, and ntC2, respectively) (Fig. 6c, d, Supplementary Fig. 14m-o). Comparison of highly expressed genes between subclusters from different datasets within each ntC extracted a common gene set including *Sox9* and *Scx* (Fig. 6e, Supplementary Data. 7). In particular, comparison within ntC0 and ntC3 corresponding to tenogenic lineage population^25^ showed 468 commonly expressed genes including *Egr1*, *Tgfb2*, *Mgp*, and *Thbs2* (Fig. 6e, f), which are upregulated during tendon development^25,26^. On the other hand, cardiac NCCs did not express definite tenocyte markers *Tnmd* and *Mkx* (Supplementary Data. 7), indicating that *Sox9*^high^/*Scx*^high^ NCCs are likely distinct from cells in the tenocyte lineage.

When the top 7 OCs were listed according to the number of parent (hub) TFs belonging to the above 468 genes, *Scx*-containing OC69 and *Sox9*/*Scx*-containing OC86 were included whereas another *Scx*-containing network OC58 was not (Supplementary Fig. 14p, Supplementary Data. 7), indicating the existence of similarities and differences in *Scx*-involving GRNs between cardiac NCCs and tenocyte progenitors. Moreover, *Egr1*-containing OC4 and OC33 were also included in the top 7 OCs (Supplementary Data. 7). Among 49 ‘child’ genes of *Egr1*, 25 genes (51%) were found in the above 468 genes (Supplementary Data. 7), indicating that *Sox9*^+^/*Scx*^+^ cardiac NCCs and tendon progenitors share *Egr1* as a key TF.

### Comparison of GRNs between *Sox9*^+^/*Scx*^+^ NCCs and cartilaginous cells

Because *Sox9*^+^/*Scx*^+^ progenitor cells give rise to *Sox9*^+^/*Scx*^−^ chondrocytes in addition to *Sox9*^−^ /*Scx*^+^ tenocytes/ligamentocytes^24^, we compared OC levels between the Visium clusters (evCs and lvCs) containing *Sox9*/*Scx*-enriched NCCs and cartilaginous cells (Fig. 7, Supplementary Fig. 15). evC4 and lvC31 represented the endocardial cushion and valvular tissues enriched for *Sox9*^+^/*Scx*^+^ NCCs and exhibited OC profiles similar to clusters representing tracheal cartilaginous tissues (eVC8 and lvC25, respectively) (Fig. 7a-d, Supplementary Fig. 15a, b). All of these clusters shared OC4, OC86 and OC91 (Fig. 7e-g, i-k). By contrast, OC58 (containing *Scx, Meox1, Sall3, Sox5, Sox6* and *Snai2*) was prevalent only in *Sox9*^+^/*Scx*^+^ NCC- containing clusters (Fig. 7h, l, Supplementary Fig. 9n). Among TFs belonging to OC58, *Meox1* and *Sall3* were highly expressed in the cushion mesenchyme, but only very weakly in the trachea (Fig. 7m-r, Supplementary Fig. 16a-t). Furthermore, *Meox1* and *Sall3* expression was downregulated in NCC-derived chondrocytes of the mandibular arch (Supplementary Fig. 17a, b, Supplementary Data 8), as evidenced by investigation of published datasets^27^. iPS-derived tendon progenitors did not express *Sall3* either, but express *Meox1* (Supplementary Fig. 14l) ^25^. These results indicate that cardiac NCCs and developing tenocytes and chondrocytes share a common set of GRNs, whereas some components of transcription networks are specific for cardiac NCCs, which may play important roles in cell fate determination and differentiation to suppress skeletogenic programs.

**Fig. 7.**
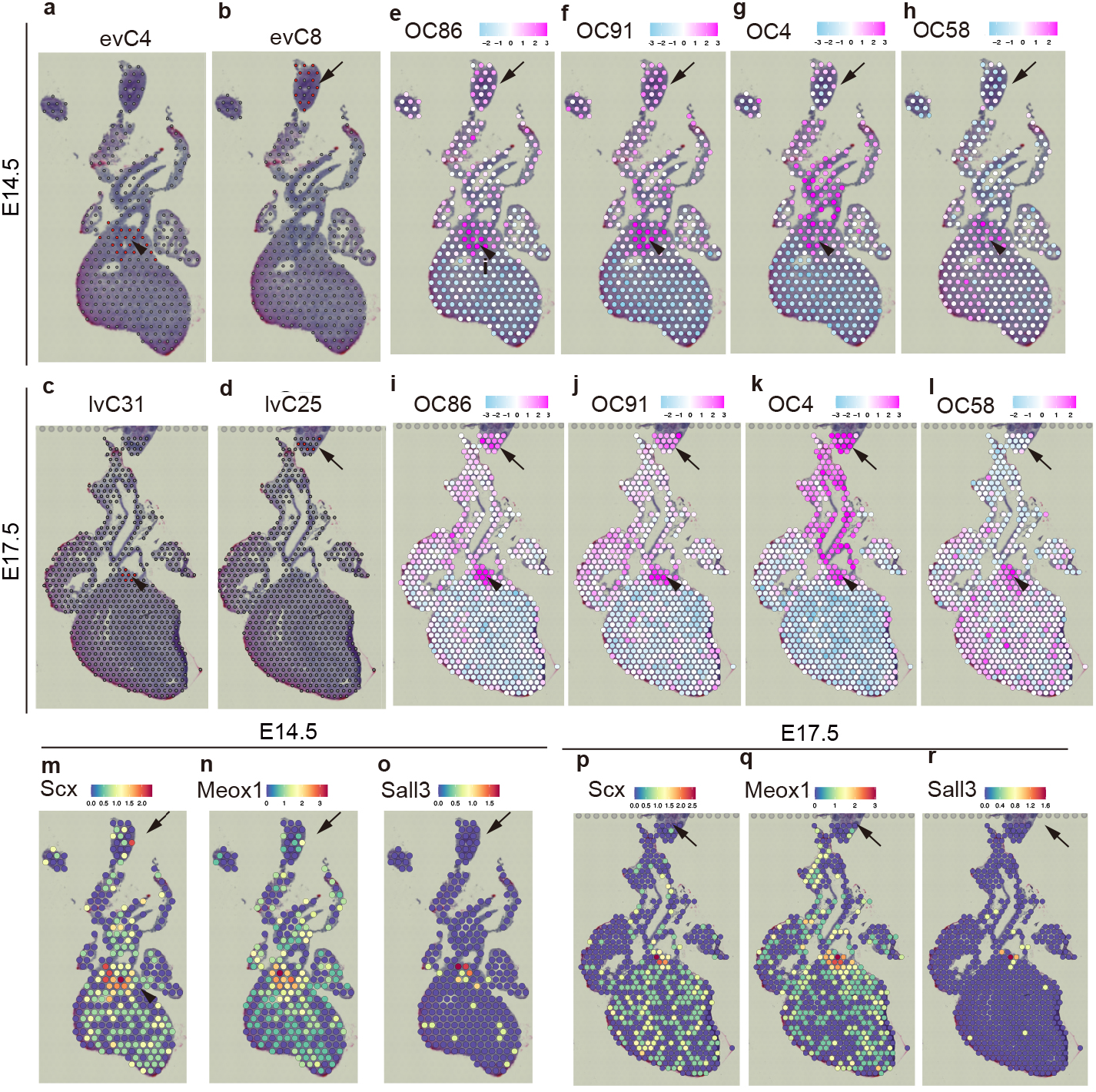
Similarity and difference in GRNs between cardiac cushion and tracheal tissues. (a-d) Cluster distribution in the Visium plots at E14.5 (a, b) and E17.5 (c, d). (e-l) Spatial overall community (OC) expression patterns in the Visium plots at E14.5 (e-h) and E17.5 (i-l). (m-r) Spatial expression patterns of key TFs in OC58 at E14.5 (m-o) and E17.5 (p-r). Arrowheads, cardiac cushion tissue; arrows, tracheal tissue.

### Difference between cardiac NCCs and skeletogenic NCCs in the pharyngeal arches

Many of cardiac NCCs and skeletogenic NCCs in the pharyngeal arches share the same origin from preotic rhombomeres^8^. To test whether they also shared the same ectomesenchymal progenitors in the pharyngeal arches, we genetically traced the cell lineage using *Dlx1-CreERT2;R26-EYFP* mice, which can label ectomesenchymal NCCs in the first and second pharyngeal arches at E9.5 (Fig. 8a). When tamoxifen was administered at this stage, EYFP-positive *Dlx1*-lineage cells were mainly observed in the anterior pharyngeal arch region (Fig. 8a). In the cardiac outflow tract region, EYFP-positive cells were sporadically detected, which were also positive for Sox9 and SM22α (Fig. 8b, c). In addition, EYFP-positive cells were found in the smooth muscle layers of the pharyngeal arch arteries, great vessels and coronary arteries, and in the aorticopulmonary septum (Fig. 8d-g). Only a small number of EYFP-positive *Dlx1*-lineage cells in the developing heart indicates that many cardiac NCCs are destined for the cardiac outflow tract before regional identities in the pharyngeal arches are specified by the *Dlx-*code and that even NCCs with the *Dlx*-code appear to differentiate differently from those in the pharyngeal arches.

**Fig. 8.**
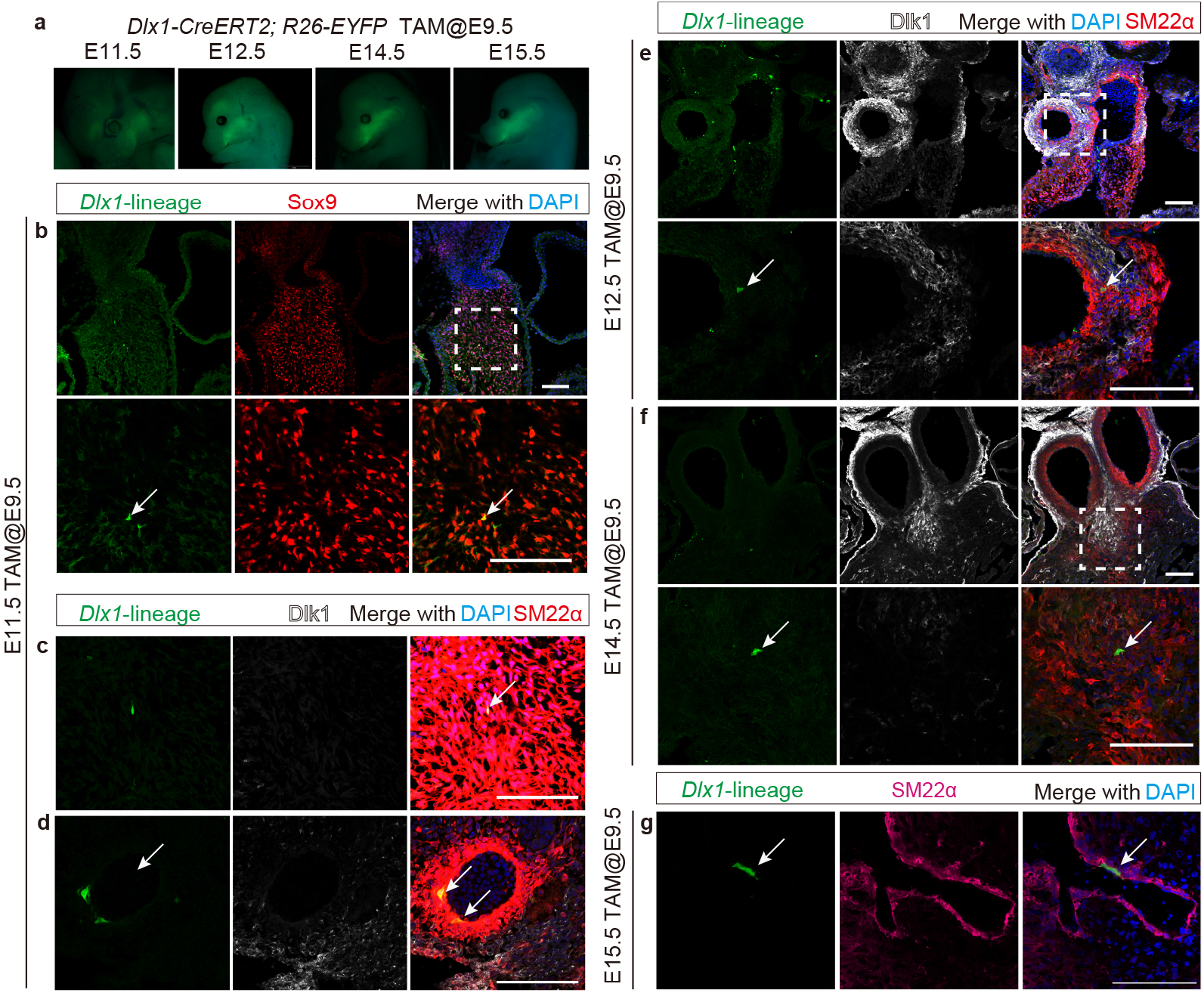
*Dlx1*-lineage cells migrating into the cardiac outflow tract. (a) Lineage tracing of Dlx1-positive cells in the craniofacial region of *Dlx1-CreERT2;R26-EYFP* mice treated with tamoxifen at E9.5. (b-g) Immunohistochemistry of EYFP (*Dlx1*-lineage) with co-staining for Sox9 (b), SM22α (c, d, g) or Dlk1 (e, f) at E11.5 (b-d), E12.5 (e), E14.5 (f) and E15.5 (g). Boxed areas are magnified in lower panels. Arrows indicate *Dlx1*-lineage cells. Nuclei are stained with DAPI. Scale bars, 50 μm.

## Discussion

In this study, we demonstrate cardiac NCC lineage diversity with inference of GRNs involved in differentiation processes based on single-cell multiome analysis (Fig. 9). Cardiac NCCs were divided into two major populations, smooth muscle(-like) and non-muscle mesenchymal cells, and one minor Schwann cell progenitor/melanoblast-like cell population. Smooth muscle(-like) and non-muscle mesenchymal lineages are derived from immature cushion-forming NCCs through a *Sox9^+^*/*Scx^+^*teno-chondrogenic progenitor-like stage. Comparison to developing chondrocytes indicates cardiac NCC-specific components of GRNs, which may suppress skeletogenic programs.

**Fig. 9.**
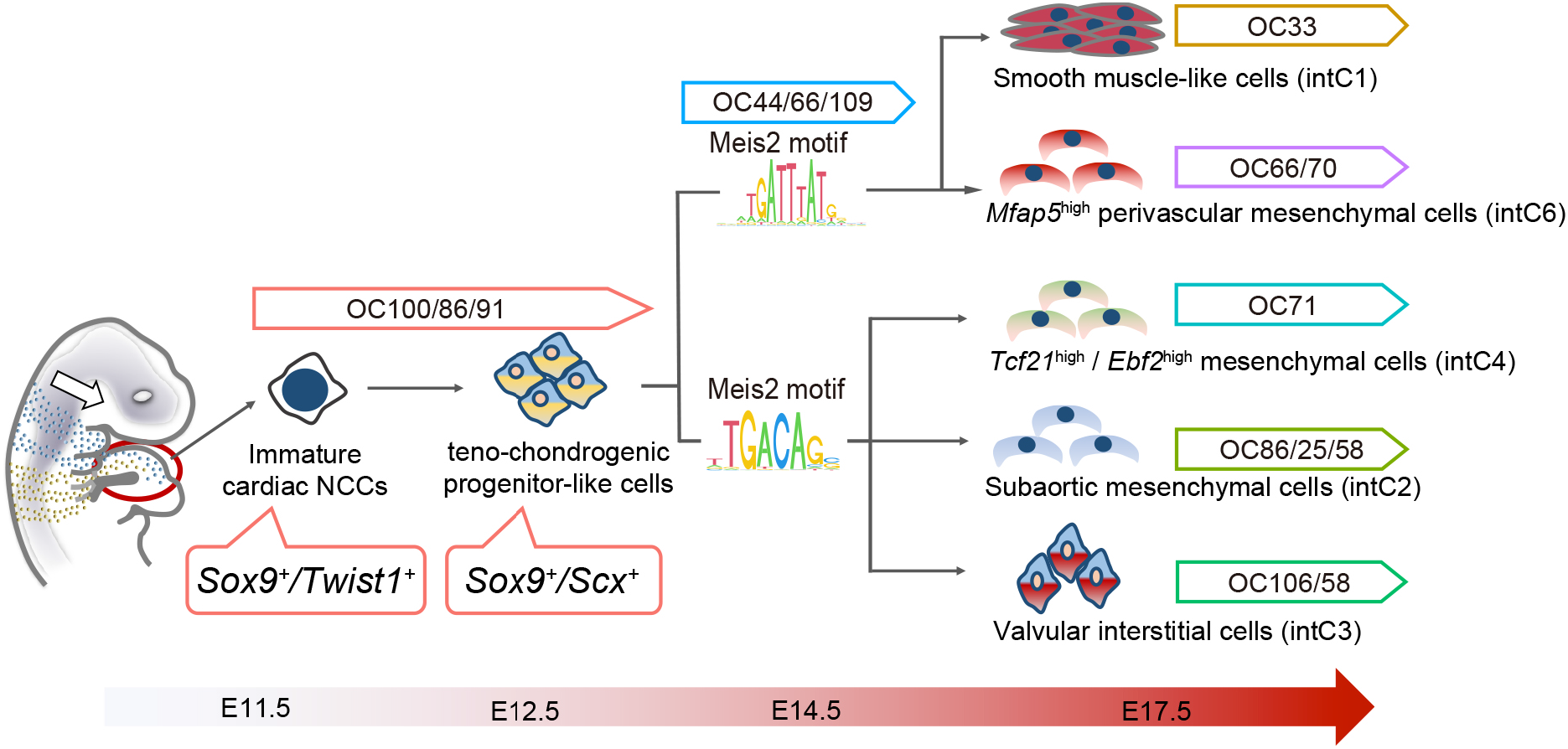
Scheme of GRNs underlying cardiac NCC lineage diversity.

Recently, Chen and colleagues reported single-cell transcriptomic landscape of cardiac NCCs, proposing the developmental path from NCCs migrating into the heart to mesenchymal and vascular smooth muscle lineages^15^. The NCC subpopulations and lineage relationship inferred in the present study are largely consistent with those reported by Chen et al., but there are some notable differences. First, they showed that the cluster representing valvular interstitial cells was characterized by *Tcf21* expression. But *Tcf21* was highly expressed in intC4 as well as immature mesenchymal and valvular interstitial clusters intC0 and intC3. Unlike intC0, intC3 and Chen’s valvular interstitial cluster c11^15^, intC4 was also characterized by *Ebf2* and allocated to tissues surrounding the great vessels in the 10x Visium platform, indicating a distinct cell population. Given that Ebf2 is a key TF regulating brown adipocyte development^28,29^, intC4 may be a mesenchymal cell population containing precursors of periaortic brown adipocytes, which were recently identified as of NC origin^30^. Secondly, Chen et al. designated the *Penk*-enriched cluster c9 as a cushion mesenchymal population involved in the formation of the aorticopulmonary septum^15^. By contrast, the present analysis found *Penk* expression broadly in smooth muscle(-like) and mesenchymal clusters, especially enriched in intC5, which bridged between the immature mesenchymal cluster intC0 and smooth muscle(-like) cluster intC1 (Supplementary Fig. 4h). Interestingly, genes characterizing a subpopulation of Chen’s c9 (sc0) such as *Lrp1b, Gabrb2*, and *Syt1* (Fig. 1l) were also enriched in intC5, indicating that this subpopulation may represent smooth muscle precursors. This inference is also supported by the connectivity of GRNs based on the multimodal analysis and Cre-mediated genetic lineage tracing. Finally, we could not find direct evidence supporting the transition from NCC-derived pericytes to microvascular smooth muscle cells reported by Chen et al., probably because they detected this transition by subclustering the mural cell population at postnatal stage P7, which was not included in the present study.

The present study further inferred GRNs involved in divergent cardiac NCC differentiation by integrating two different algorithms on independent datasets; SCENIC-based nonparametric Bayesian network analysis on the Fluidigm C1 scRNA-seq dataset and multimodal *cis*-regulatory network analysis based on the RNA-seq and ATAC-seq data from the 10x Genomics scMultiome platform. This approach enabled us to robustly extract hierarchical GRNs underlying differentiation of immature mesenchymal NCCs into smooth muscle(-like) and non-muscle mesenchymal lineages. Immature mesenchymal NCCs were characterized by GRNs containing *Meis2*, *Sox9* and *Scx*. Consistently, genetic lineage tracing using different Cre-expressing mouse lines revealed that most of cardiac NCC-derivatives differentiate through a *Sox9^+^*/*Scx^+^* intermediate state, reminiscent of *Sox9^+^*/*Scx^+^* teno-chondrogenic progenitors. *Sox9-*containing GRNs and Sox9-binding motifs remained enriched in non-muscle mesenchymal lineages, especially in the subaortic mesenchymal cluster intC2, consistent with the expression of Col2a1, a downstream target of Sox9^23^, in the NCC-composed mesenchymal condensation. By contrast, *Sox9* and its binding motifs were downregulated along smooth muscle differentiation with concomitant upregulation of *Osr1*, which has been reported to suppress *Sox9* expression^19^. Thus, Osr1 may suppress Sox9-dependent genetic programs to activate smooth muscle differentiation programs involving its downstream gene *Foxc1*. Interestingly, the present analysis inferred that both *Sox9* and *Osr1* were downstream of Meis2 through different binding motifs, which were differently enriched in smooth muscle(-like) and non-muscle mesenchymal lineages, suggesting a possibility that switching of different Meis2-bindig motifs may be involved in smooth muscle(-like)/mesenchymal bifurcation.

*Sox9^+^*/*Scx^+^* teno-chondrogenic progenitor-like NCCs were clustered in the roots of semilunar valves as well as in the subaortic mesenchymal condensation. In analogy to the tendon-bone attachment unit formed by *Sox9^+^*/*Scx^+^*progenitors^31,32^, *Sox9^+^*/*Scx^+^*NCCs in the roots of semilunar valves may constitute an attachment unit between the cushion tissue and valvular leaflets. Indeed, the present 10x Visium analysis indicated that cardiac NCCs and developing chondrocytes shared a common set of GRNs involving *Egr1*, a critical TF for tendon maturation and matrix deposition^33^. On the other hand, the analysis also revealed that some components of transcription networks, such as *Sall3,* were specific for cardiac NCCs. Unlike skeletogenic cranial NCCs, cardiac NCCs give rise to neither bone nor cartilage in mice and humans, indicating the existence of inhibitory mechanisms preventing activation of skeletogenic programs. Indeed, even NCCs with the *Dlx*-code did not exhibit osteo-chondrogenic differentiation. Previous reports have demonstrated that knockout of *Ets1*, *fibronectin 1* (*Fn1*) or *a disintegrin and metalloprotease 19* (*Adam19*) in mice results in NCC-derived cartilage formation in the heart^34–36^. It would be interesting in the future to explore the possibility of corporation between the cardiac NCC-specific transcription networks and these factors.

It is noteworthy that the coronary artery and aortic valves, to which are highly contributed by cardiac NCCs, are prone to calcification, but the developmental basis for this propensity is not fully understood. The similarities and differences in GRNs between cardiac NCCs and skeletogenic progenitors may provide a clue to understanding the pathogenesis of calcification. Thus, the present finding will provide a basis for understanding the role of NCCs in cardiac development and some insights into studies on the cell fate determination and differentiation as well as various issues in stem cell biology from basic and clinical viewpoints.

## Methods

### Animals

*Wnt1-Cre*^13^, *Rosa26-loxp-stop-loxp-EYFP* (*R26-EYFP*)^14^, *Sox9-EGFP*^37^, *Sox9-CreERT2*^38^, *Scx-CreL*^31^, *R26-CAG-loxP-stop-loxP-tdTomato* (*Ai14*)^39^ and *Dlx1-CreERT2*^40^ mice have been described previously. *ScxTomato* transgenic mice were generated by injecting the ∼11 kb *Scx*-*tdTomato* transgene into pronuclei of fertilized eggs (Shukunami C, unpublished). Mutant mice were maintained on a mixed C57BL/6J × ICR background. All animal experiments were approved by the Ethics Committee for Animal Experiments of The University of Tokyo and the Committee of Animal Experimentation of Hiroshima University. Mice were housed at 23±2°C with a relative humidity of 50-60% and light cycles with 12 hours light and 12 hours dark.

### Single-cell RNA sequencing (scRNA-seq) of cardiac NCCs

Cardiac NCCs labeled by *Wnt1-Cre; R26-EYFP* mice at E11.5, 12.5, 14.5 and 17.5 around sub-region of aorticopulmonary septum were dissociated by using 0.25 w/v% trypsin / 1 mmol/L EDTA·4Na solution (Wako) at 37 °C, 15 minutes. Enzymes were neutralized with equal volume of DMEM (Wako) with 10% fetal bovine serum (FBS) (SIGMA). Cell suspensions were filtered through a 35 μm nylon mesh cell strainer(FALCON 352235). Cells were kept on ice until cell sorting. Cell sorting was performed in a fluorescence-activated cell sorting (FACS) AriaⅡcell sorter (BD Biosciences). EYFP-positive single cells were captured on the Fluidigm C1 Medium-cell (10-17 μm cell diameter) integrated fluidic circuit (IFC) for single-cell cDNA libraries for mRNA sequencing. EYFP-positive cells were loaded on the IFC chips at concentration of 300 cells per μL. All captured cells were photographed to check whether cells were truly EYFP-positive and were dissociated for single cells by using KEYENCE BZ-X710. Single-cell cDNAs were prepared on Fluidigm C1 system by using SMARTer v4 Ultra Input Low RNA kit for the Fluidigm C1 System (Clontech). Only single-cell cDNAs confirmed by imaging data were performed quality check (QC) by using Agilent 2200 TapeStation system and quantified by Qubit (Thermo Fisher). High quality cDNAs were further constructed sequencing libraries by using Nextera XT DNA Sample Preparation Kit (Illumina). Each single-cell cDNAs were sequenced 50 pair-end on the Illumina HiSeq 2500.

### Single-cell Multiome (scRNA-seq and scATAC-seq) of cardiac NCCs

Cardiac NCCs labeled by *Wnt1-Cre; R26-EYFP* mice at E14.5 and E17.5 were prepared for single-cell analysis as the same as above and cryopreserved for stock. After the preparation of cell stock, the cells were thawed in DMEM with 10% FBS and the nuclei were isolated as described in the protocol, Nuclei Isolation for Single Cell Multiome ATAC + Gene Expression Sequencing with the option of Low cell input nuclei isolation (10x Genomics). Isolated nuclei were inserted by the transposase with the fragments in the open chromatin regions for scATAC-seq. After the transposition, the nuclei were subjected to the droplet in the oil with the Gel Bead-in-emulsion (GEM) system (10x Genomics). In this process, each nucleus was performed the reverse transcription in the droplet for scRNA-seq. Each cDNA and genome DNA derived from the same cell was prepared for the construction for gene expression (GEX) and ATAC sequence library as described Chromium Next GEM Single Cell Multiome ATAC + Gene Expression (10x Genomics). Each of GEX and ATAC library was sequenced on Illumina NovaSeq6000 as manufacturer provided.

### Fluidigm C1 single-cell RNA-seq data analysis

Data analysis was performed in R version 4.1.1 and Python 3.8. Sequence output fastq files were aligned to indexed mouse (*Mus musculus*) genome reference (GRCm38/mm10) with extra EYFP DNA sequences by using HISAT2 software version 2.1.0. Gene expression counts were calculated with the featureCounts function from the Rsubread version 1.34.7 package using R. All scRNA-seq data have 388 cells, including 45 cells at E11.5, 100 cells at E12.5, 117 cells at E14.5 and 126 cells at E17.5. The 309 cells mapped at least 1 EYFP sequence were used for further analysis, including 39 cells at E11.5, 81 cells at E12.5, 79 cells at E14.5 and 110 cells at E17.5.

At first, Each of count data was converted into a Seurat object which contains 15124 features (genes) and 309 samples (cells) by using Seurat version 3.2.0 R package^41^. The percentage of mitochondrial genes against detected genes per cell was calculated by using the top 200 genes associated with mitochondria, Mouse MitoCarta 2.0 (BROAD INSTITUTE). 294 cells which contain the number of feature RNA > 2000 and the percentage of mitochondrial genes < 0.1 were used for further analysis, including 34 cells at E11.5, 78 cells at E12.5, 76 cells at E14.5, and 106 cells at E17.5. The cell cycle score was calculated by using the CellCycleScoring function in the Seurat package. This cell cycle score was used for normalization by using the SCTransform function to reduce the batch effect of cell cycle. All single-cell data were subjected to clustering through the Louvain algorithm based on a k-nearest neighbor graph on the Euclidean distance in PCA space and were projected onto the two-dimensional UMAP space. Each differentially expressed gene (DEG) corresponding to each cluster was calculated by using FindAllMarkers function. The parameters were set as default. The statistical significance was performed with a Wilcoxon Rank Sum test and the threshold was limited by log fold change was more than 0.25.

Second, RNA velocity analysis was performed by using velocyto^43^, including velocyto.py version 0.17.17 Python package and velocyto.R version 0.6. The expression counts of each spliced and unspliced genes were calculated through mapping to mouse genome reference (GRCm38/mm10) by using run_smartseq2 function from velocyto.py. Then, RunVelocity and show.velocity.on.embedding.cor function in SeuratWrappers version 0.1.0 package in R were used for RNA velocity analysis.

### Single-cell multiome (scRNA-seq and scATAC-seq) data analysis

Sequence output fastq files were aligned to mouse (*Mus musculus*) genome reference (refdata-cellranger-arc-mm10-2020-A-2.0.0) provided by 10x Genomics to produce gene expression counts by scRNA-seq and fragments by scATAC-seq in Cell Ranger ARC v2.0.0 pipeline (10x Genomics). Then, gene expression counts and fragments were converted into Seurat objects which contained 32285 features and 593 cells and 344 cells at E14.5 and E17.5, respectively by using Seurat version 4.0.4 and Signac version 1.0.4 R package^43,44^. These gene expression data were merged into a dataset, whereby overlapping peaks across samples were merged to reduce batch effect. The percentage of mitochondrial genes was calculated by the PercentageFeatureSet function. The dataset was filtered for the number of feature RNA > 2000, the number of feature RNA < 7500, the number of count RNA > 3000, the number of count RNA < 45000, the number of feature ATAC > 1000, the number of feature ATAC < 45000, the number of count ATAC > 3000, the number of count ATAC < 100000, TSS enrichment > 2, nucleosome_signal < 4 and percentage of mitochondrial genes < 15. After filtering, the dataset contained 32285 features and 434 cells (E14.5 was 236 cells and E17.5 was 198 cells). Cell cycle scoring and normalization were performed as described above. Then, each of the RNA- and ATAC-seq datasets was subjected to dimensional reduction into a two-dimensional UMAP space. These datasets were analyzed by using weighted nearest neighbor (WNN) analysis for clustering based on the integration of both information^43^. DEGs and peaks corresponding to each cluster in the RNA- and ATAC-seq dataset, respectively, were estimated by using wilcoxauc.Seurat function provided by the presto R package with default parameters. TF binding motifs in highly accessible chromatin regions were analyzed by ChromVAR^20^. For the integration of Fluidigm and 10x Genomics scRNA-seq data, canonical correlation analysis (CCA)-based integration method was used to reduce technical batch in Seurat R package^40^.

We created an R program under Seurat and Signac R packages to construct *cis*-regulatory TF network from scMultiome data (scRNA-sed and scATAC-seq). This R program also predicts and visualizes which were enhancer- or silence-like peaks. This program includes several steps. Step0; Filtering TFs with logFC > 0.25, pct_in > 25, and avgExpr > 0.25 among scMultiome clusters. Step1; TF and target sets with mean normalized SCT greater than 0.25 for any cluster are subjected to analysis. Step2; scATAC-seq data are used for searching *cis*-co-accessible networks (CCANs) with Cicero version 1.3.4.11 R package^45^. Detected CCANs are used to assign each accessible peak to a gene by linking to the closest transcription start sites (TSSs). TSS-linked peaks with normalized values greater than 0.25 for any one of the clusters were used for further analysis. Step3; Extracted peaks correlated with TSS through CCANs are matched with TF binding motifs by motifmatchr version 1.14.0 R package^46^ registered as vertebrates in JASPAR2020^47^. Step4; Matched peaks are filtered with positive or negative correlation for detecting enhancer- or silencer-like regions, respectively, with some thresholds; correlation score between peaks and TSS (named as scorethreshold, default is 0.1) and spearman correlation score between peaks and gene expressions (named as theshold_cor, default is 0.05). Step5; these motifs are filtered by highly conserved sequences among Euarchontoglires with phastCon score^48^. We used the conservation score bigwig file based on the mm10, mm10.60way.phastCons60wayEuarchontoGlire.bw, to import R with rtracklayer package. An average conserved score with more than 0.25 was used for the visualization. In this study, to compare between smooth muscle-like (SML; mC4, mC5, mC6_0, mC6_1, and mC7) and non-muscle mesenchymal population (NML; mC0, mC1, mC2, mC3, and mC9), genome accessible average peak data with threshold (logFC > 0.2 and avgExpr > 0.2) was used for conservation score filtering step.

### Estimation of GRNs

We used SCENIC^17^ version 1.1.2.2 R package to extract the gene combinations inferred by gene regulatory motifs and SiGN-BN NNSR version 0.16.6^16^ to construct GRNs based above SCENIC gene combinations in the super-computing software provided by Human Genome Center, the Institute of Medical Science, the University of Tokyo. The Fluidigm C1 scRNA-seq expression count data were converted into transcripts per million (TPM) value to reduce the effect of mapping bias because of the characteristics of full-length RNA-seq. TPM data without fC11, including 15124 genes and 284 cells was analyzed in SCENIC algorithm. Gene filtering was performed default geneFiltering function in SCENIC. Then, Spearman correlation was calculated by using runCorrelation function and the combination of genes, the absolute value of correlation value > 0.03 (default), were used for further analysis. The data extracted by only positive correlation > 0.03 were used for the calculation of TFs – gene co-expression modules by using runGenie3 function. Furthermore, the modules were purified by using RcisTarget to get regulons which contain each TF binding motif among the TF-gene combinations. The input 2 mouse gene regulatory motif databases for SECNIC were mm9-500bp-upstream-7species.mc9nr.feather and mm9-tss-centered-10kb-7species.mc9nr.feather files. We used motifEnrichment_selfMotifs parameter set to “not” which were detected in the only TFs in databases but all motifEnrichment because of enabling detection of minor TFs–genes combinations. It might include a broad TF – gene combination so we run the SiGN-BN NNSR algorithm to construct a GRN based on nonparametric Bayesian estimation on the SHIROKANE supercomputer at the University of Tokyo. The data run on SiGN-BN NNSR were TPM expression data filtered by only genes, above Seurat FindAllMarkers function result set the parameter of only.pos = FALSE, and except cells labelled fC11. SiGN-BN NNSR was run at the following parameter, m; the maximum number of parents that each gene can have was set to 1000, T; the number of iterations of the subnetwork estimation by the neighbor node sampling and repeat algorithm was set to 1000000, skel-type TXT-- skel; the restriction of the combination between Parent and Child edge was set to above regulons result analyzed by SCENIC filtered by Genie3Weight > 0.004.

### Analysis of GRN community

The TF-TF subnetworks were extracted in the result of GRN. TF-TF subnetworks were performed community detection with some overlapping to summarize the relationship in TF nodes by using linkcomm R package^18^. After the community detection, the score of each community per cell in the scRNA-seq data was calculated by some steps. Step1; each gene of TPM expression count per community was converted into z-scores to reduce the gap of the kinds of genes of the community. Step2; each averaged gene expression score was averaged with some genes belonging to each community per cell. Step3; Z-scored gene expression scores were averaged in every cluster to calculate the score of each community per cluster. The spatial community detection was also estimated similarly. The scores of the community were visualized as heatmap with pheatmap R package. Genes belonging to OCs in which they were connected to TF(s) through the first edge were used for gene ontology analysis with Metascape^42^.

### Spatial transcriptome of embryonic hearts

Embryonic hearts labeled by *Wnt1-Cre; R26-EYFP* mice at E14.5 and 17.5 were embedded in a chilled OCT compound and immediately frozen on the metal plate in a liquid nitrogen bath. Frozen samples were cut at the thickness of 10 μm in the cryostat and placed on the Visium Spatial Tissue Optimization Slide to confirm the time to lyse sections and on the Visium Spatial Gene Expression Slide to get RNA-seq data. The lysis time to extract RNA from the section was determined by the results of Visium Tissue Optimization Kits (10x Genomics). Then, the cryosections on the Visium Spatial Gene Expression Slide were stained with Hematoxylin and Eosin (HE) and captured images with KEYENCE BZ-X710. After imaging, the sections were lysed in 24 and 18 minutes at E14.5 and 17.5, respectively, with Visium Spatial Gene Expression Reagent Kits (10x Genomics). After that, mRNAs on the slides were synthesized for cDNA and constructed for the sequencing library as the manufacturer provided. Each of the libraries was sequenced on Illumina NovaSeq6000 as manufacturer provided.

### Spatial transcriptome data analysis

Total sequence output fastq files (E14.5_1, E14.5_2, E17.5_1, E17.5_2) were aligned to indexed mouse (*Mus musculus*) genome reference (GRCm38/mm10) with extra EYFP DNA sequences as same as above and gene expression counts were also calculated by using Space Ranger version 1.1.0. The gene expression counts were converted into Seurat objects and extracted only the space of interest to exclude the noise. After normalization with the SCTransform function, the spatial gene expression counts data containing 15076 features and 702 spots in merged E14.5 data and 15729 features and 1428 spots in merged E17.5 data. Each spatial gene expression dataset was performed dimensional reduction in two-dimensional UMAP space and spot clustering as above. DEGs across cell clusters were calculated by using FindAllMarkers function. The parameters were set as default. The statistical significance was performed with a Wilcoxon Rank Sum test and the threshold was limited by log fold change was more than 0.25.

For the detection of cardiac NCCs on the space, the spatial gene expression counts mapped at least 1 EYFP sequence were used for further analysis. To characterize spatial information of Fluidigm C1 scRNA-seq clusters, the spatial gene expression and single-cell gene expression data were integrated with the label transfer method in Seurat R package^41^. The prediction of spatial identity of single cells was visualized on HE-stained images.

### Analysis of public datasets

We used the public scRNA-seq dataset of mouse tenogenic differentiation from iPSCs^25^ to compare cNCCs scRNA-seq data. The count matrix data from T0 to T9 was downloaded from GEO under accession number GSE168451. Ensemble ID was converted to gene id by BioTools.fr (https://www.biotools.fr/mouse/ensembl_symbol_converter). Read counts for duplicated gene id were added and summarized. Expression matrix data were converted into a Seurat object and analyzed as above. However, the filtering conditions are as follows, nFeature_RNA > 1500 & nFeature_RNA < 6000 & percent.mt < 20. To integrate this data with Fluidigm C1 cNCCs scRNA-seq data, the CCA method was used as above. To avoid the reduction of genes because of the compression with integration, DEGs in iPS data were calculated for iPS clusters 0 and 3 corresponding to undifferentiated T0 while DEGs in sC data were calculated for immature sC3. Venn diagrams were visualized with ggvenn R package.

We also used the public scRNA-seq dataset of mouse embryonic craniofacial tissues for searching osteocytes/chondrocytes gene expression patterns^27^. The count matrix data at E14.5 was downloaded from The FaceBase data repository (Record ID 1-DTK2, doi: 10.25550/1-DTK2; http://facebase.org). These data were converted into a Seurat object and analyzed as above. However, the filtering conditions are as follows, nFeature_RNA > 2000 & nFeature_RNA < 6000 & percent.mt < 5.

### Lineage tracing experiment

For lineage tracing of Sox9-positive cells in *Sox9-CreERT2; R26-tdTomato* mice at E11.5, the pregnant female mice were injected with 2.5 mg of 4-hydroxy tamoxifen (Sigma-Aldrich). For lineage tracing of Dlx1-positive cells in *Dlx1-CreERT2; R26-EYFP* mice at E9.5, the pregnant female mice were induced oral administration of tamoxifen (Sigma-Aldrich) in corn oil (Sigma-Aldrich) at a dose of 0.1mg/g body weight. These embryos were sampled at appropriate stages.

### Immunohistochemistry

Embryos were fixed in 4% paraformaldehyde phosphate buffer solution (Nacalai Tesque) for 3 hours at 4°C. The fixed embryos were embedded in OCT compound (Sakura Finetek) through stepwise sucrose substitution and stocked at –20°C until sectioning. These samples were cut by using cryostat at thickness of 10 μm. For immunohistochemistry, frozen sections were rinsed with phosphate-buffered saline (PBS) and induced transparency processing with 0.3 % Triton-X100 in PBS. Blocking was used 3% bovine serum albumin (BSA) in PBS. Primary antibodies mixed with 3% BSA buffer were on the sections for overnight at 4°C. After washing with PBS, secondary antibodies mixed with 3% BSA buffer were on the sections for 2 hours at room temperature. Antibodies were below, SM22α (ab14106, Abcam, 1:800), GFP (04404-26, Nacalai Tesque, 1:1000), GFP (600-101-215, Rockland, 1:500), Collagen typeⅡ (600-401-104-0.1, Rockland, 1:400), smooth muscle myosin heavy chain 11 (ab53219, Abcam, 1:200), Jagged 1 (AF599, R&D systems, 1:500), Dlk1 (AF1144, R&D systems, 1:500), Sox9 (A19710, ABclonal, 1:500) and Alexa Fluor (488, 555 and 647)-conjugated secondary antibodies (Abcam, 1:200). Nuclei was stained by 4’,6-diamidino-2-phenylindole dihydrochloride (DAPI) solution. Immunofluorescence images were captured by a Nikon C2 confocal microscope and BZ-X810.

### Data availability

The Fluidigm C1 scRNA-seq data have been deposited in the Gene Expression Omnibus under accession code GSE201417. The 10x Genomics scMultiome data and Visium spatial transcriptome data have been deposited in DDBJ Sequence Read Archive (DRA) under accession code DRA012897 and DRA010734, respectively. Public mouse tenogenic differentiation from iPSCs scRNA-seq data was downloaded from GEO under accession number GSE168451. Public craniofacial scRNA-seq data was downloaded from the FaceBase data repository (Record ID 1-DTK2, doi: 10.25550/1-DTK2; http://facebase.org).

### Code availability

Scripts used for Fluidigm C1 scRNA-seq, 10x Genomics scMultiome, Visium analysis, and public data analysis are available at https://github.com/iaki-dev/Iwase_etal_2022.

**Source data**

## Acknowledgements

We thank Yoshinori Tamada (Hirosaki University) for technical guidance and the use of SiGN-BN NNSR on the super-computing resource provided by Human Genome Center (the University of Tokyo), Kiyomi Imamura and Erina Ishikawa for technical assistance of sequence. A.I. was a doctoral student fellow of Fostering Advanced Human Resources to Lead Green Transformation (GX)(SPRING GX)in the University of Tokyo. This work was supported by Core Research for Evolutional Science and Technology (CREST) of the Japan Science and Technology Agency (JST), Japan (JPMJCR13W2), Grant-in-Aid for the Japan Society for the Promotion of Science (JSPS) KAKENHI grant numbers 19H01048, 21K19519, 22H04991 (to H.K.), 19K08308 (to S. M.-T.), 17J11177, 20H04858, 20K15858 (to H.H.), 16H06279 (PAGS) (201141, 202112, 211046), the Mitsubishi foundation, the Fugaku Foundation (to H.K.) and RIKAKEN HD (to A.I.).

## Author contributions

A.I., Y.U., D.S., Y.K., S.M.-T., and H.K. conceived the study and designed experiments. A.I., Y.U., D.S., M.K., H.H., K.M., and A.T. performed experiments. C.S., Y.U. and Y.K. supported the experiments. A.T., S.Y., S.F., T.K., M.S., Y.S., Y.W., and H. Aburatani provided the sequence platform. A.I., Y.U., S.Y., S.F., T.K., and S.N. analyzed the sequence data. C.S. and H. Akiyama provided mutant mice. A.I. and H.K. wrote the manuscript with help from other authors.

## Materials & Correspondence

H.K. had materials and correspondence of this study. C.S. and H.A. had the mutant mice such as S*ox9-EGPP, Scx-tdTomato, Sox9-CreERT2, Scx-CreL,* and *Ai14*.

