## Supplemental Figures for "Cardiac neural crest lineage diversity and underlying gene regulatory networks revealed by multimodal analysis"

### Supplementary Figures

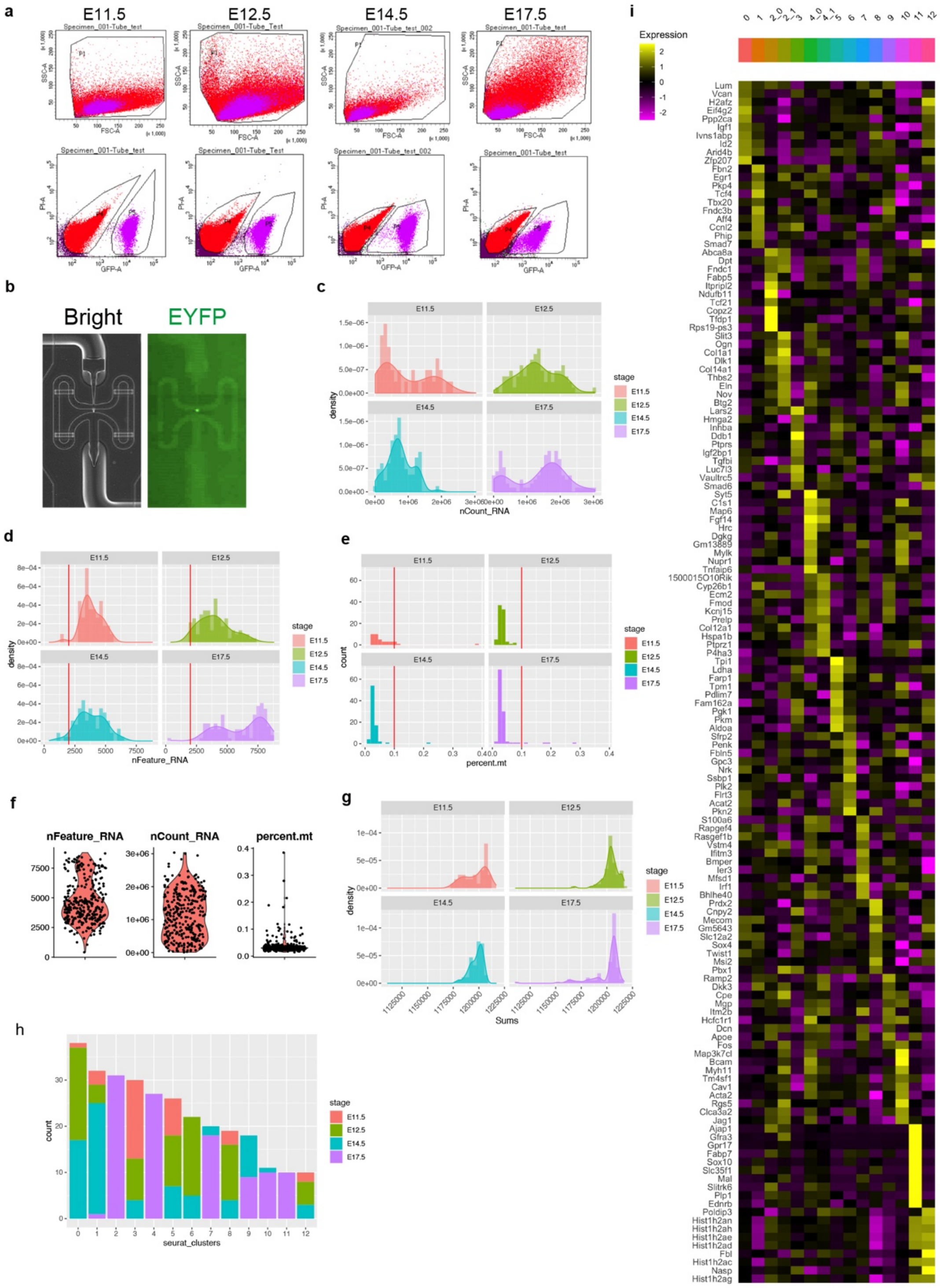


**Supplementary Figure. 1 | Quality control and overall characterization of Fluidigm C1 scRNA-seq data.**

(a) Results of the fluorescence-activated cell sorting (FACS). The upper row figures indicate the gating based on the forward scatter (FSC) and the side scatter (SSC). The lower ones indicate the gating GFP and PI to distinguish EYFP-positive (fraction P5) or -negative (fraction P4) cells.

(b) Optical and fluorescent images of a captured cell in the integrated fluidic circuit.

(c) Histograms and density plots of the number of RNA counts.

(d) Histograms and density plots of the number of detected RNA features. Red vertical lines indicate 2000, the threshold for subsequent analysis. More than 2000 RNAs were detected in 97.4% of cells (301/309).

(e) Histograms of the percentage of detected mitochondrial genes to total RNA counts. Red vertical lines indicate 0.1 %, the threshold for subsequent analysis. Less than 0.1% of mitochondrial genes to total RNA counts were detected in 96.7% of cells (299/309).

(f) Violin plots of detected counts and the percentage of mitochondrial genes among the total data.

(g) Histograms and density plots of the normalized SCT counts.

(h) Bar plots of the number of cells per cluster with the proportion of cells in different stages.

(i) Heatmap of DEGs among fCs on Fluidigm C1 scRNA-seq data.

### Supplementary Figure. 2

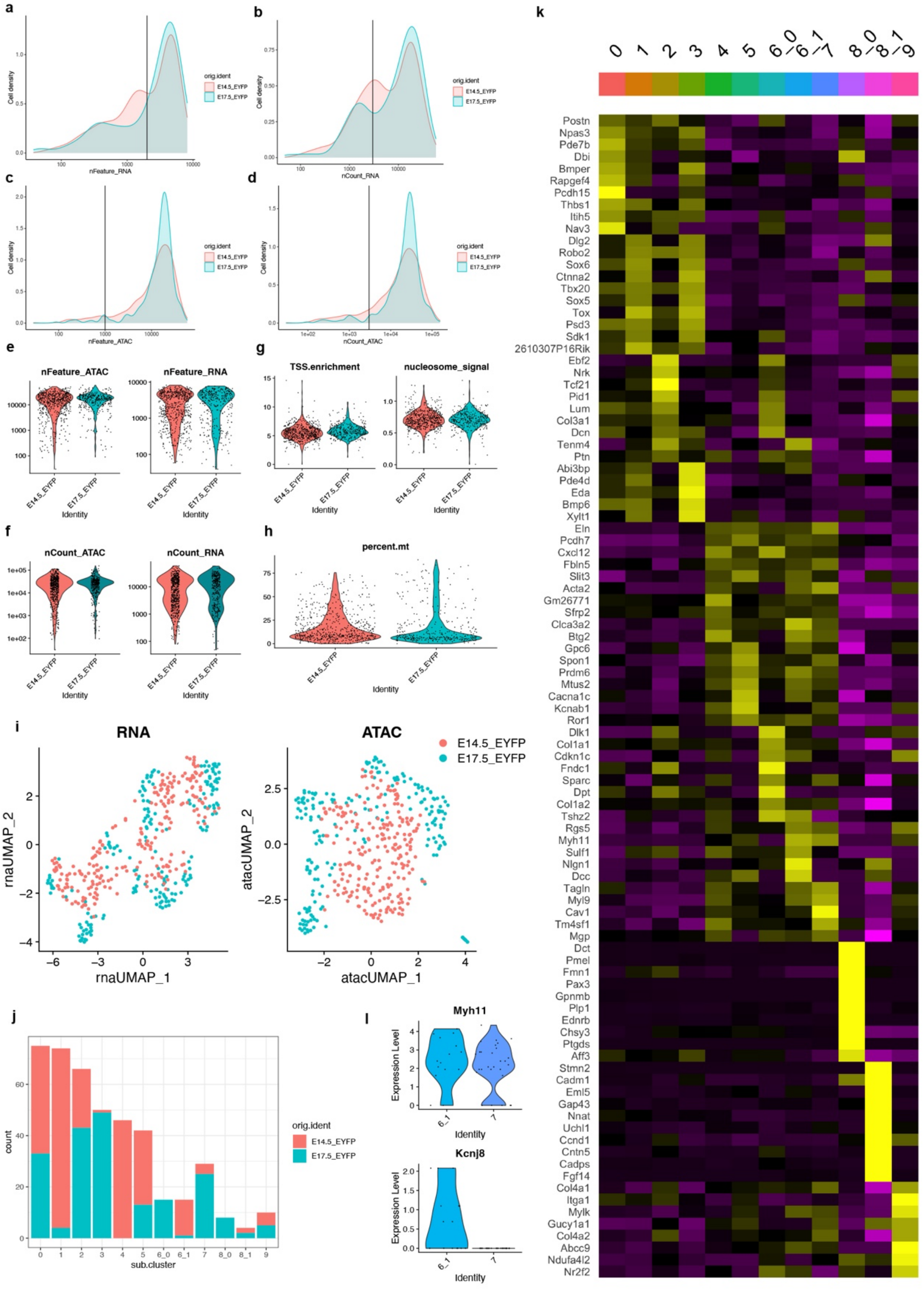


**Supplementary Figure. 2 | Quality control and overall characterization of single-cell Multiome (scRNA-seq and scATAC-seq) data.**

(a-d) Density plots of the numbers of detected RNA features (a), RNA counts (b), ATAC features (c) and ATAC counts (d) at E14.5 and E17.5. Vertical lines (a, 2000; b, 3000; c, 1000; d, 3000) indicate the thresholds for subsequent analysis.
(e-h) Violin plots of the numbers of detected RNA and ATAC (e), RNA and ATAC counts (f) and the number of transcription start site (TSS) enrichment and nucleosome signal (g) and the percentage of mitochondrial genes among total RNA counts (h) for quality control.
(i) UMAP plots of 10x Genomics scRNA-seq and scATAC-seq data.
(j) Bar plots of the number of cells per cluster with the proportion between stages.
(k) Heatmap of DEGs among fCs on 10x Genomics scRNA-seq data.
(l) Violin plots of Myh11 and Kcnj8 in smooth muscle clusters mC6_1 and mC7.

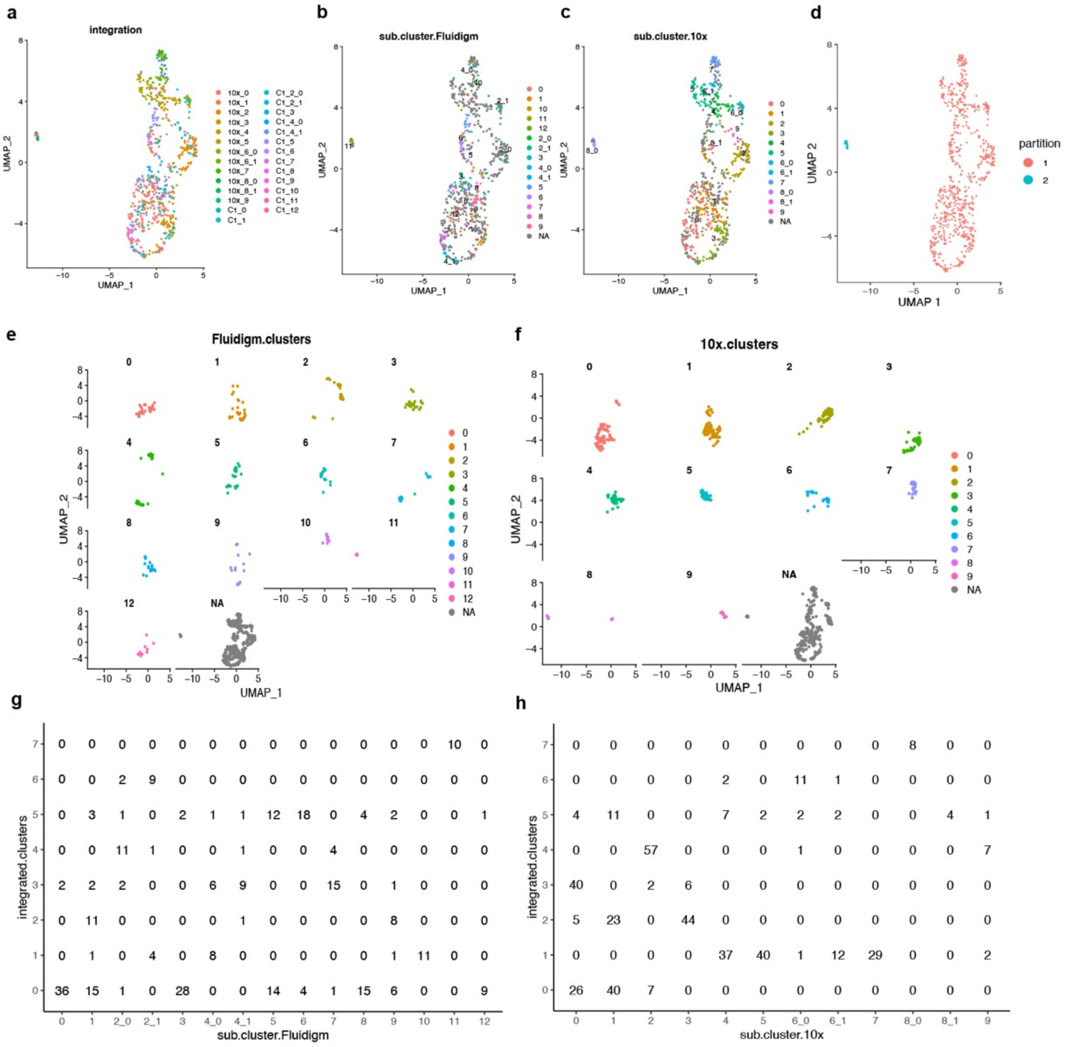


**Supplementary Figure. 3 |** Integration of Fluidigm C1 and 10x Genomics subclusters.
(a-d) UMAP plots of integrated data, colored by cluster of each platform (a), fC (b), mC (c), and partition (d). NA indicated cells of another platform.
(e-f) Split view of UMAP plots, colored by fC (e) and mC (f). NA indicated cells of another platform.
(g, h) Cross tables between fC and intC (g) and between mC and intC (h).

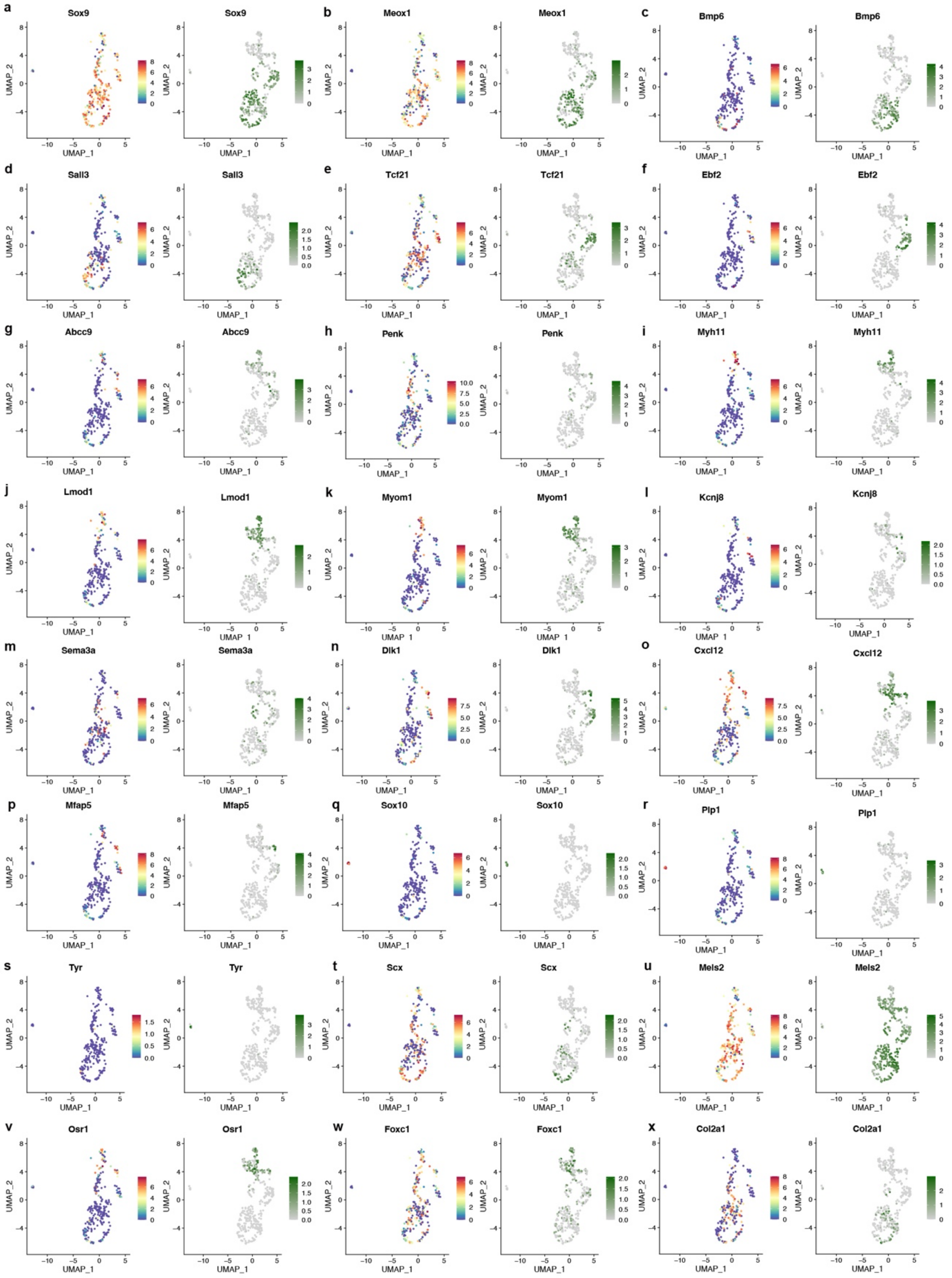


**Supplementary Figure. 4 |** Expression of marker genes on the integrated UMAP. (a-x) UMAP plots of integrated clusters were split by platforms, Fluidigm C1 scRNA-seq data (left) and 10x Genomics scRNA-seq data (right).

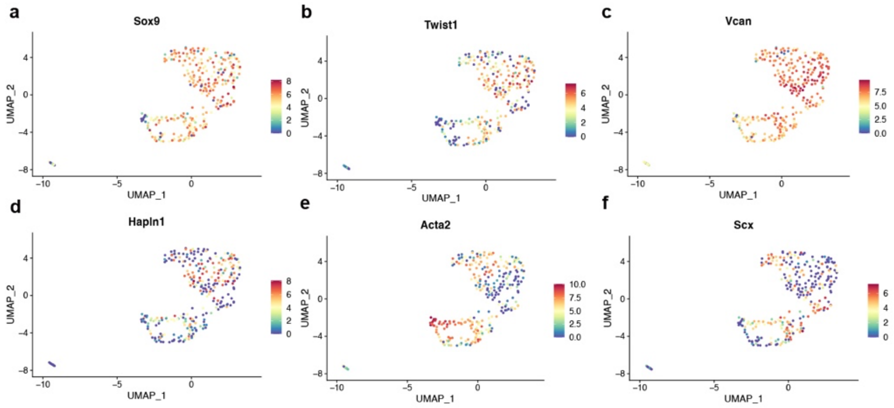


**Supplementary Figure. 5 |** Expression of marker genes on the Fluidigm scRNA-seq UMAP.

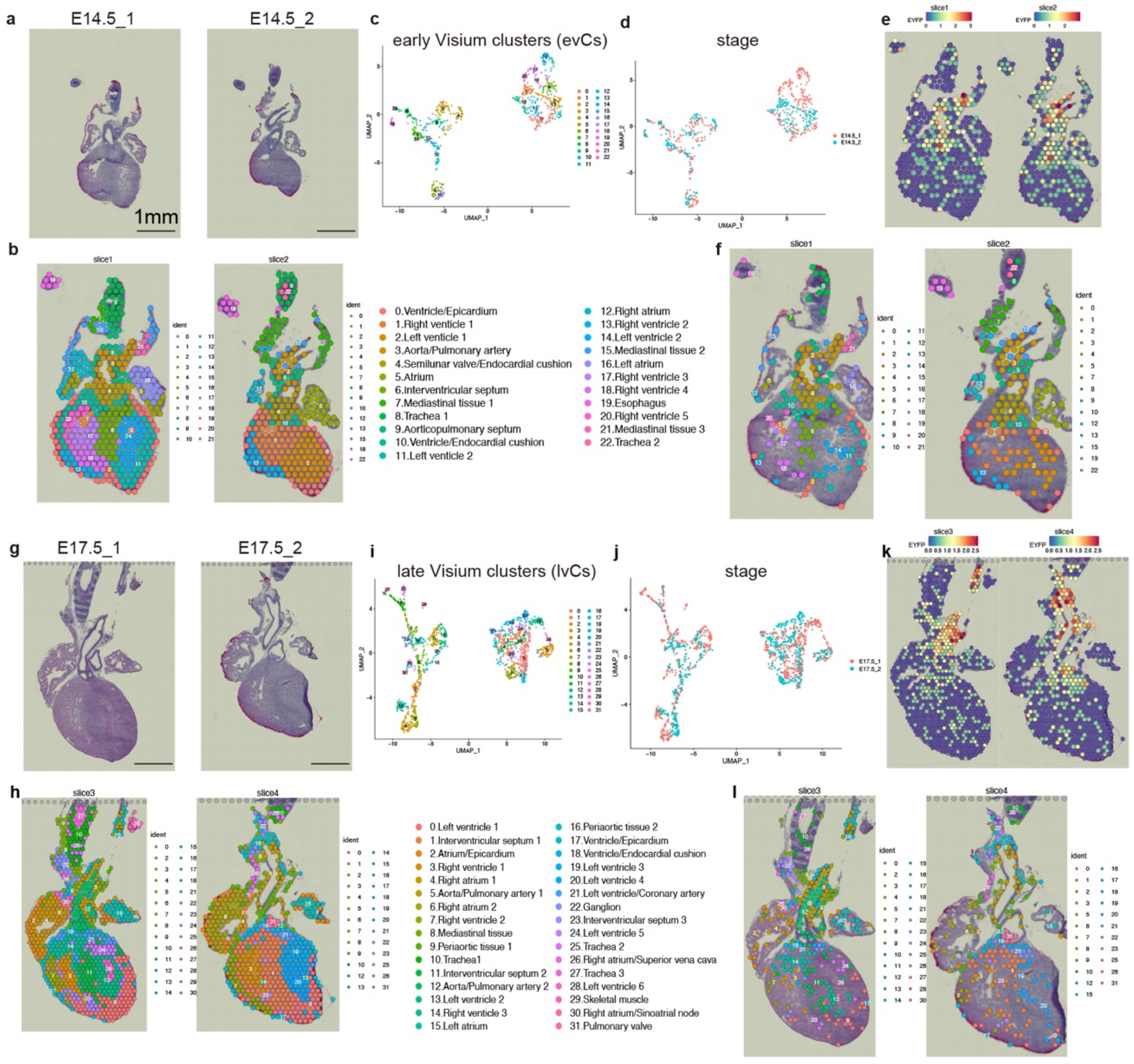


**Supplementary Figure. 6 | Spatial transcriptomics of the *Wnt1-Cre*-labeled developing heart.**

Spatial transcriptomic profiling using the 10x Visium platform at E14.5 (a-f) and E17.5 (g-l). On HE-stained images (a, g), clustering profiles are visualized (b, h) with UMAP plots, colored corresponding to early Visium clusters (evCs) (c), late Visium clusters (lvCs) (i) and stages (d, j). EYFP-expressing spots are also plotted on sections (e, k) with cluster annotations (f, l).

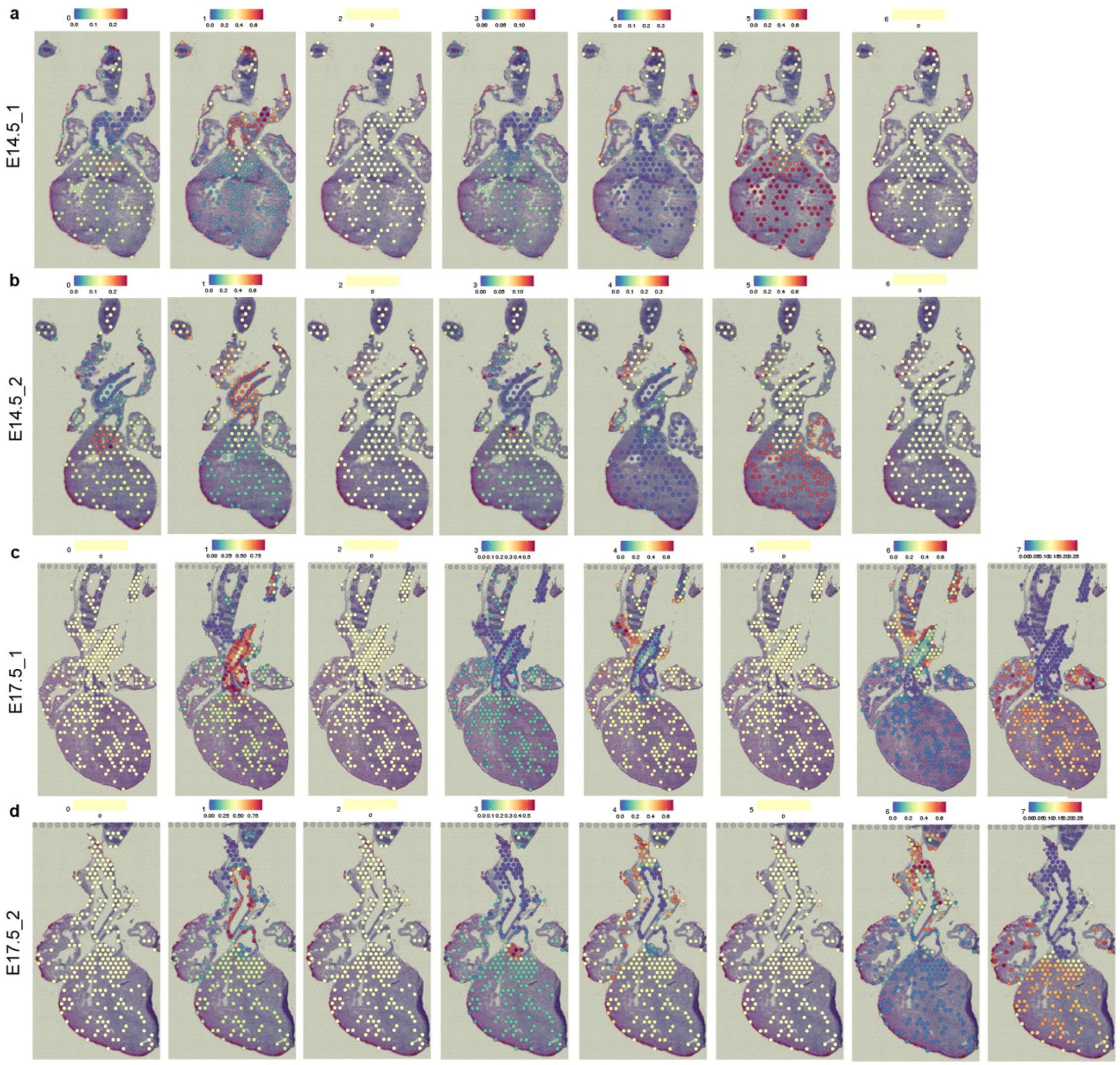


**Supplementary Figure. 7 |** Spatial identity of integrated clusters.

Prediction of integrated scRNA-seq data localization at E14.5 (a, b) and E17.5 (c, d), showing the correspondence between intCs and each spot. The color bars indicate the probability of the spatial prediction. The same stage of single-cell and spatial transcriptome data were subjected to the spatial integration analysis.

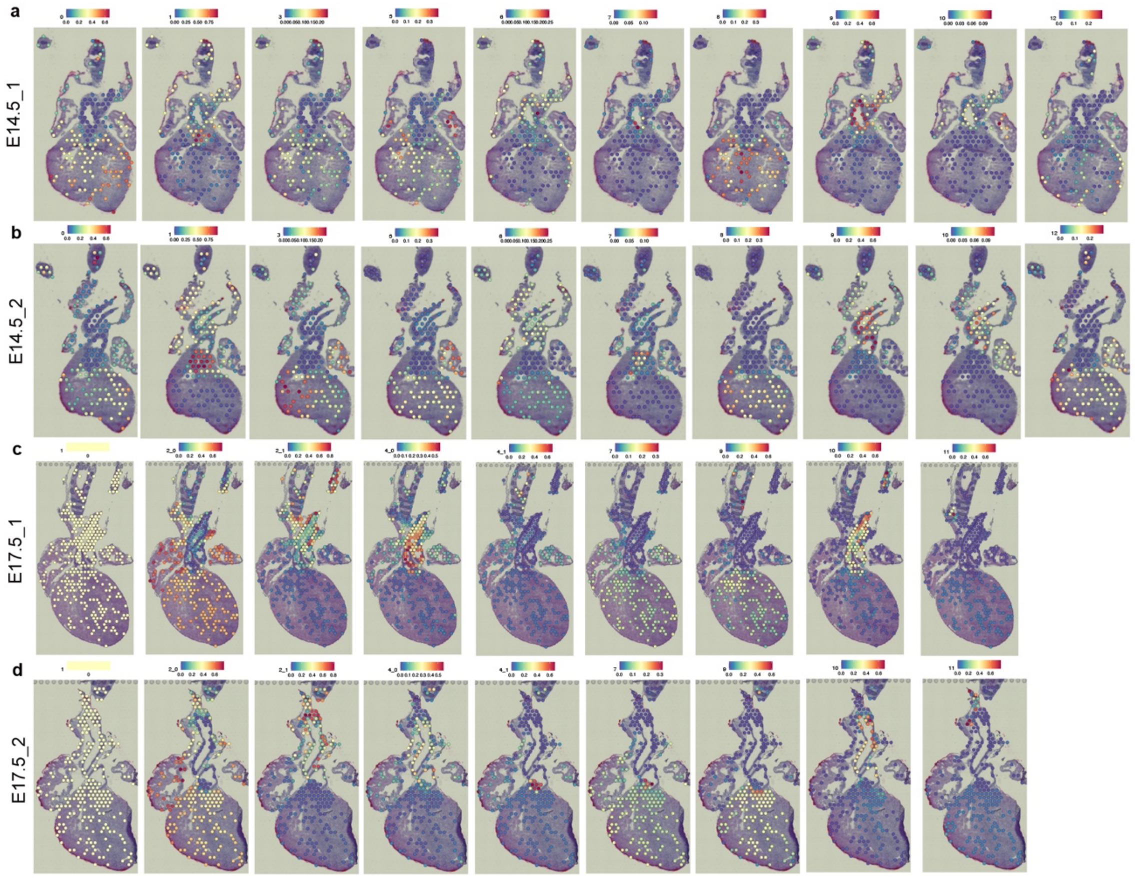

**Supplementary Figure. 8 |** Spatial identity of Fluidigm C1 single-cell clusters. Prediction of Fluidigm C1 scRNA-seq data localization at E14.5 (a, b) and E17.5 (c, d), showing the correspondence between the fCs and each spot. The color bars indicate the probability of the spatial prediction. The same stage of single-cell and spatial transcriptome data were subjected to the integration analysis.

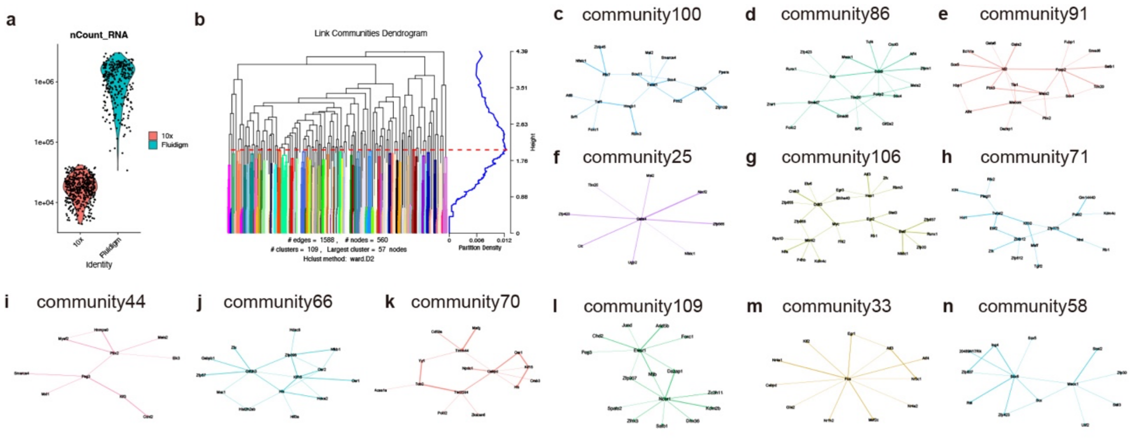


**Supplementary Figure. 9 |** Community detection of gene regulatory networks.

(a) Comparison of the number of RNA counts between 10x Genomics and Fluidigm C1 platforms.

(b) Dendrogram of community detection on the subnetwork of transcription factors based on the maximization of modularity allowed duplication. Red horizontal line indicates the threshold for the decision of communities.

(c-n) Representative community networks.

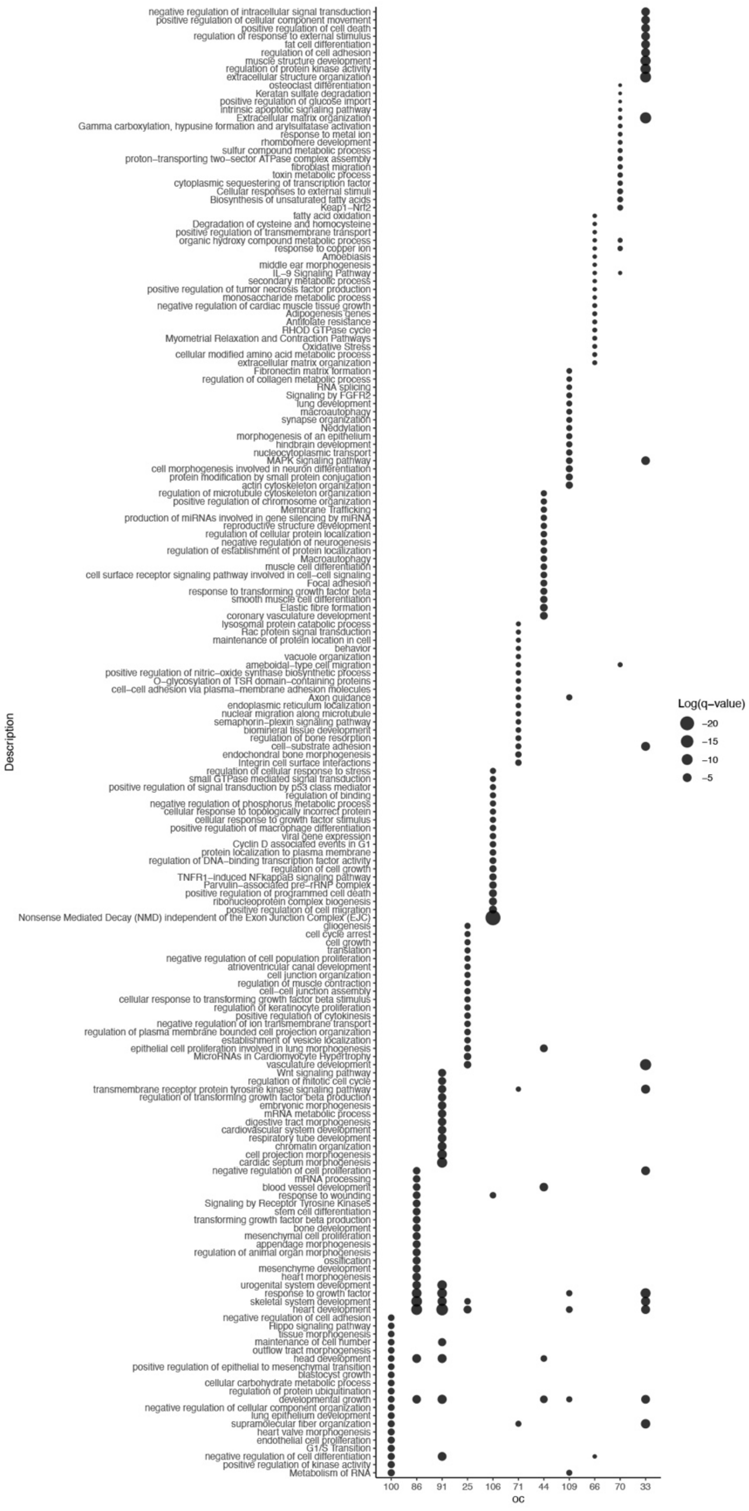


**Supplementary Figure. 10 |** Gene ontology analysis of overall community networks. The size of dot plots corresponds to log(q-value) for each term.

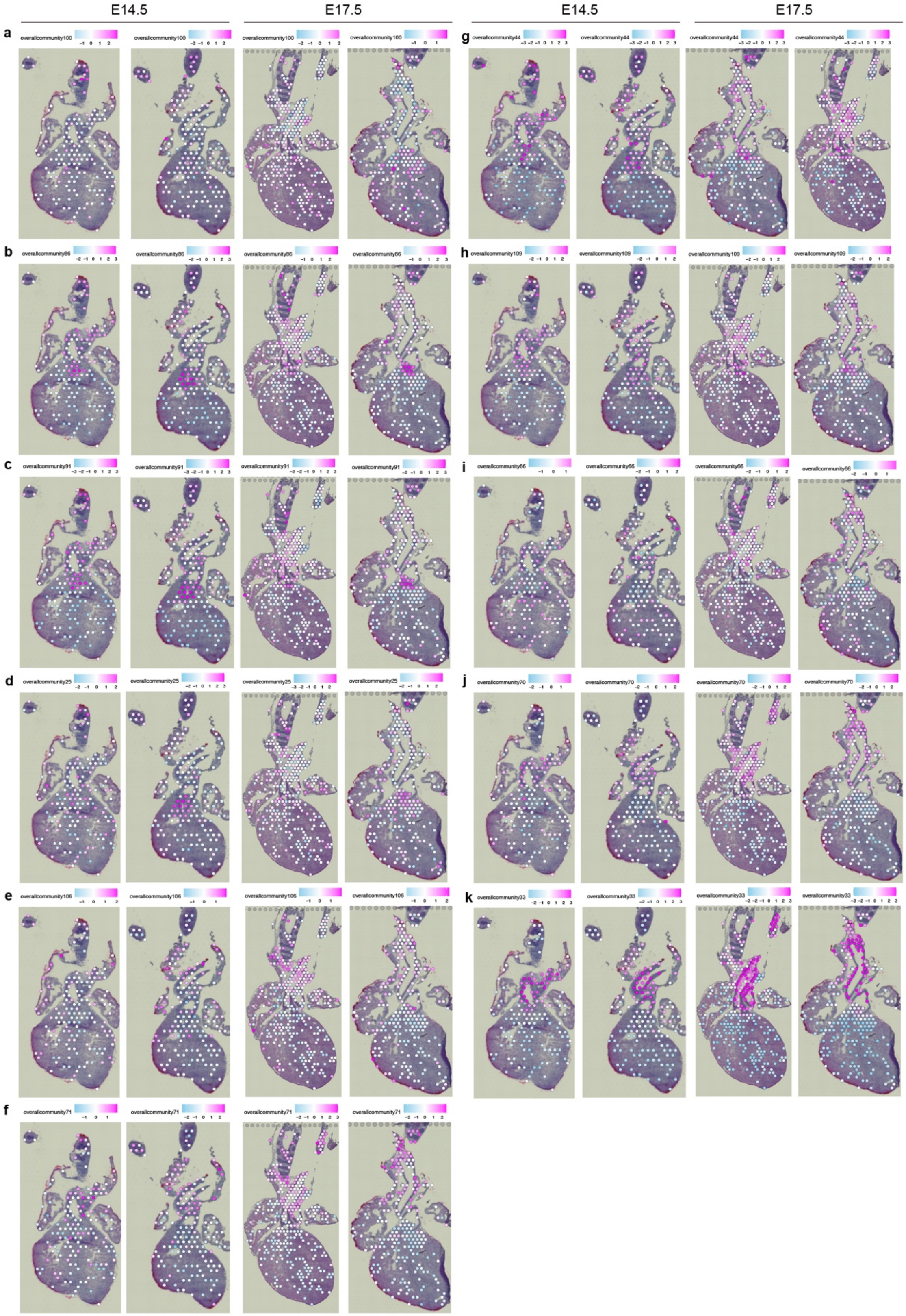


**Supplementary Figure. 11 |** Spatial overall community profiles.

(a-k) Spatial expression values of OCs were visualized on the Visium sections.

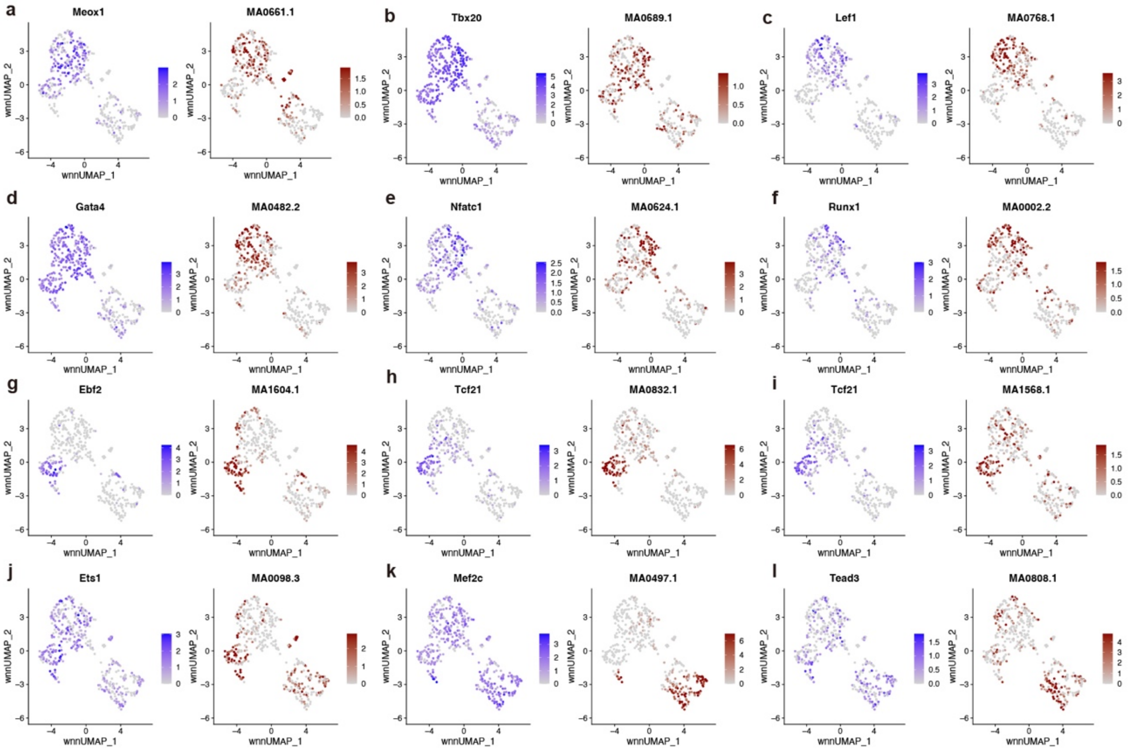

**Supplementary Figure. 12 |** Marker transcription factors identifying mC.
(a-l) RNA expression (left) and TF motif accessibility (right) were visualized on the UMAP plots with relative intensities.

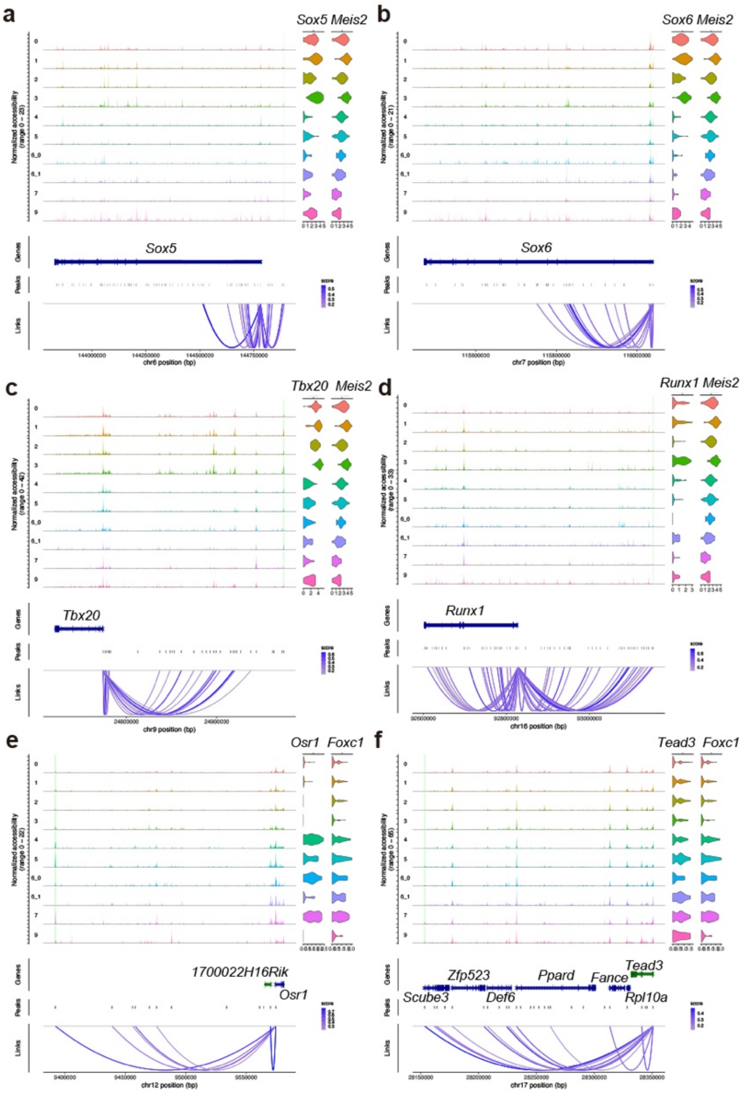


**Supplementary Figure. 13 |** Coverage plots of key transcription factors.

(a-f) Coverage plots (left) from scATAC-seq data and RNA expression of violin plots (right) from scRNA-seq data. For coverage and violin plots, each row indicates mC, with the enhancer-like region highlighted with green.

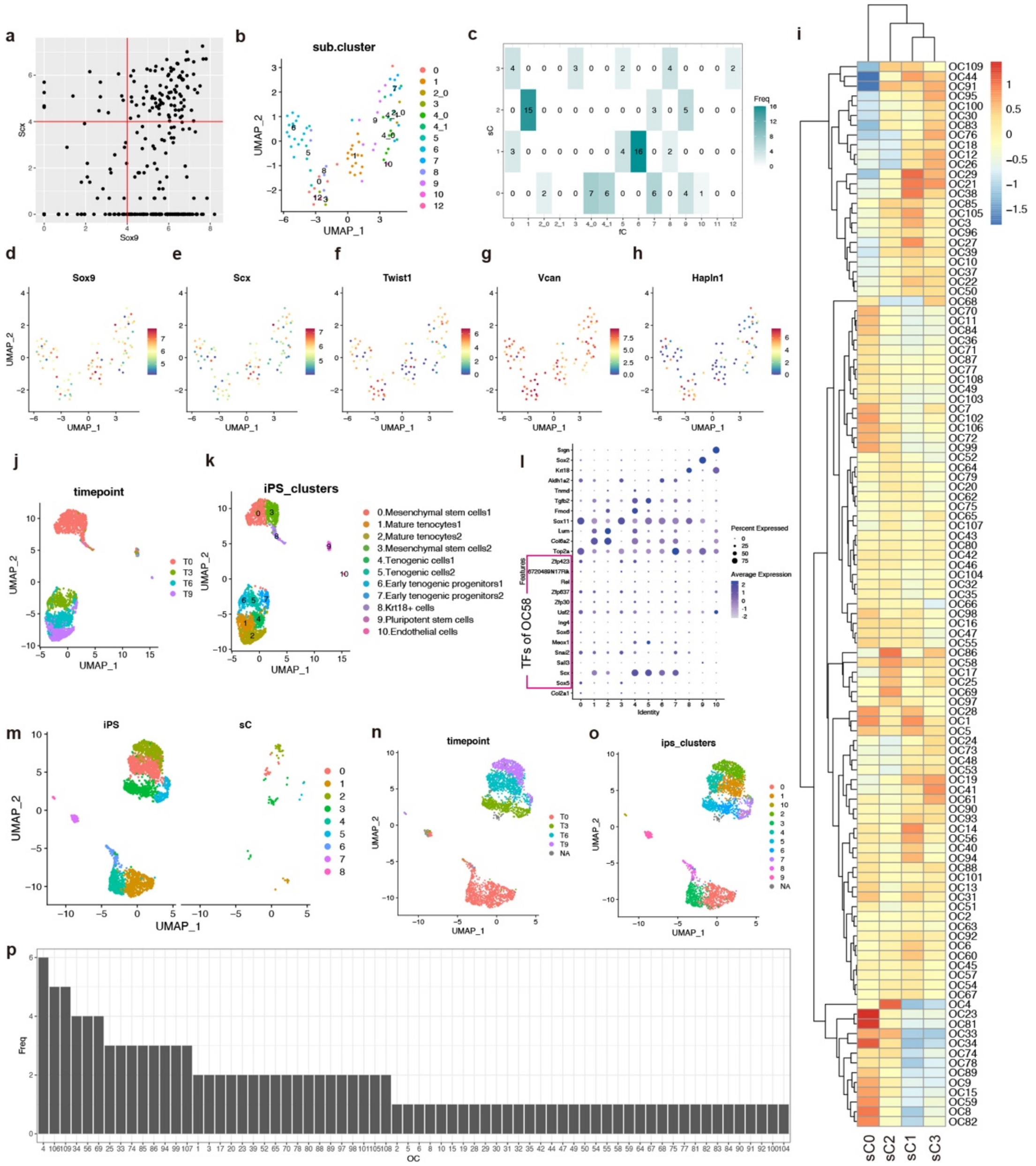


**Supplementary Figure. 14 | Comparison of** *Sox9*$^{high}$/*Scx*$^{high}$ intermediate cell population in cNCCs and tenocyte-lineage cells.

(a) Scatter plot of *Sox9* and *Scx* expression in the Fluidigm C1 data. Red vertical and horizontal lines indicate the threshold to determine *Sox9* and *Scx* high expressors.

(b) UMAP plot of *Sox9*$^{high}$/*Scx*$^{high}$ NCC subclusters (sC) labeled by fC identity.

(c) Cross table between sC and fC.

(d-h) Gene expression profies of sC UMAP plots.

(i) Heatmap of OCs among sCs on Fluidigm C1 scRNA-seq data.

(j, k) UMAP plots of iPS-derived tenocyte-lineage cell data deposited in the GEO under accession number GSE168451. Color indicates timepoint (left) and iPS clusters (right).

(l) Dot plot of marker genes and TFs of OC58 on iPS data. Each column indicates iPS cluster. Marker genes are as follows; *Top2a* (proliferative cells); *Col6a2, Lum, Tnmd* (mature tenocytes); *Sox11* (Mesenchymal stem cells); *Fmod, Tgfb2* (tenogenic cells); *Aldh1a2* (early tenogenic progenitors); *Sox2* (undifferentiated pluripotent stem cells); *Srgn* (endothelial cells).

(m-o) UMAP plots of combined scRNA-seq data from fC clusters and iPS-derivatives[25], split into sC and iPS data (m) or colored by time point (n) or original iPS cluster identity (o).

(p) Histogram of the number of parent (hub) TFs of OCs belonging to the common 468 genes shared by cNCCs and tenocyte-lineage cells.

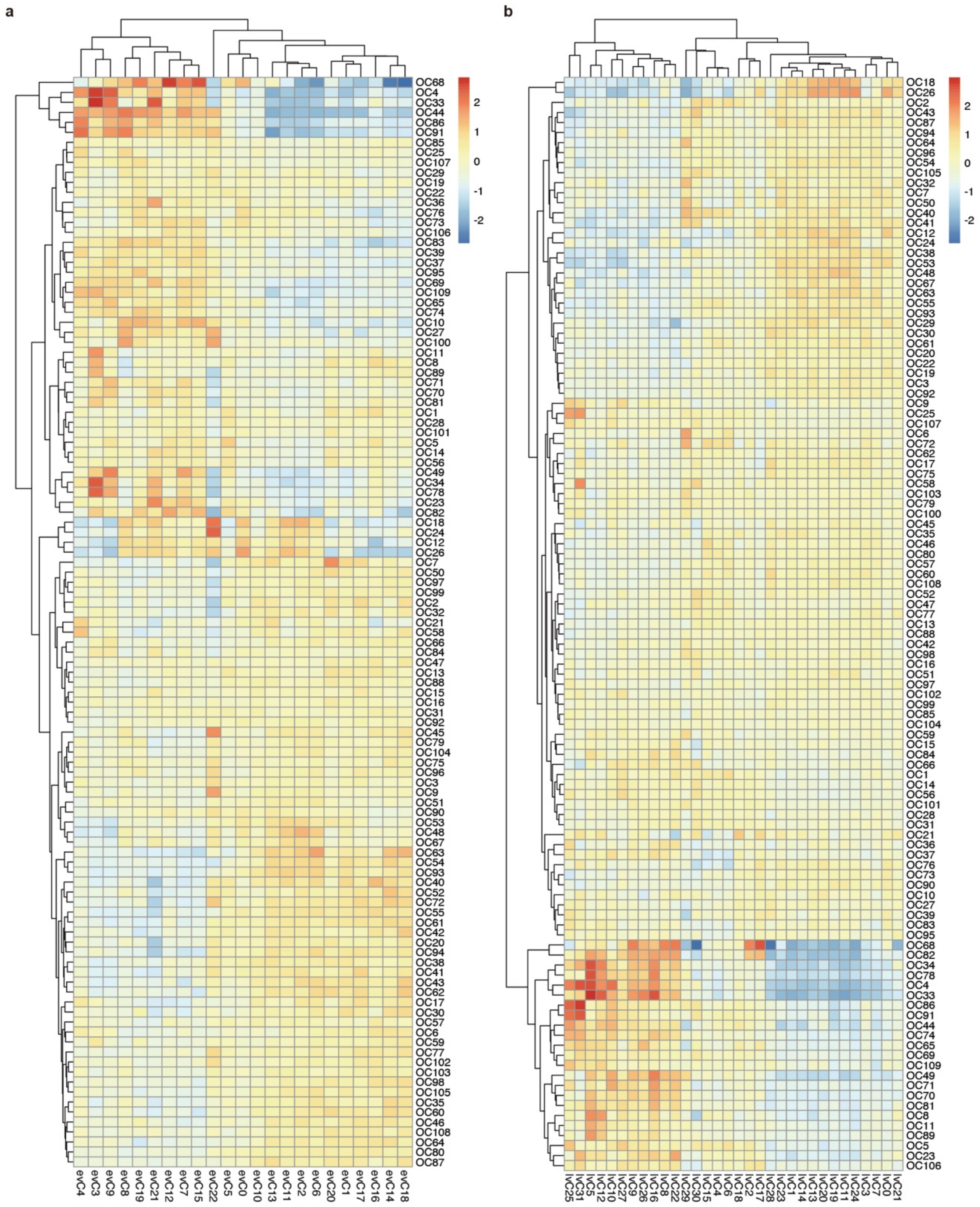


**Supplementary Figure. 15 |** Expression patterns of OCs in the Visium clusters.
(a, b) Heatmaps showing OC expression levels in E14.5 evCs (a) and E17.5 lvCs (b).

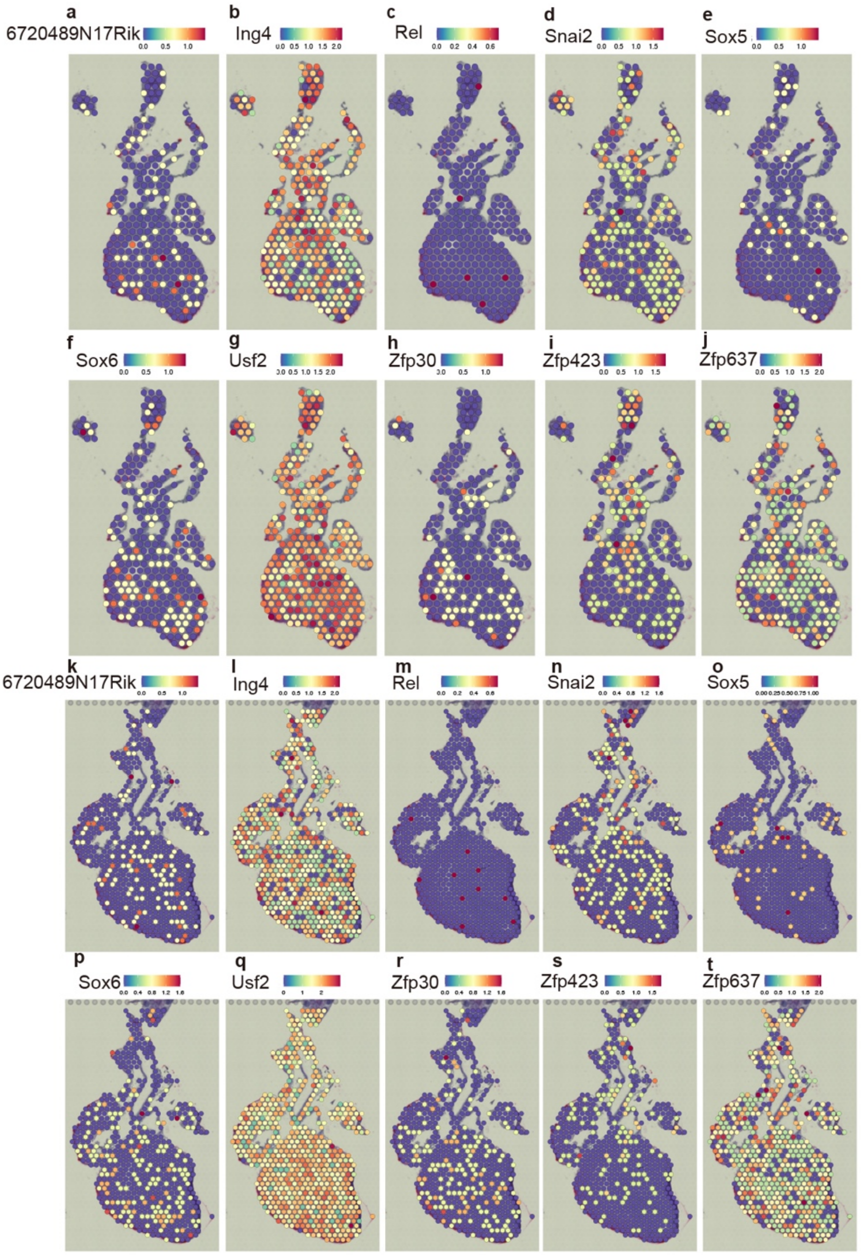


**Supplementary Figure. 16 |** Expression patterns of TFs of OC58 in Visium plots at E14.5 (a-j) and E17.5 (k-t).

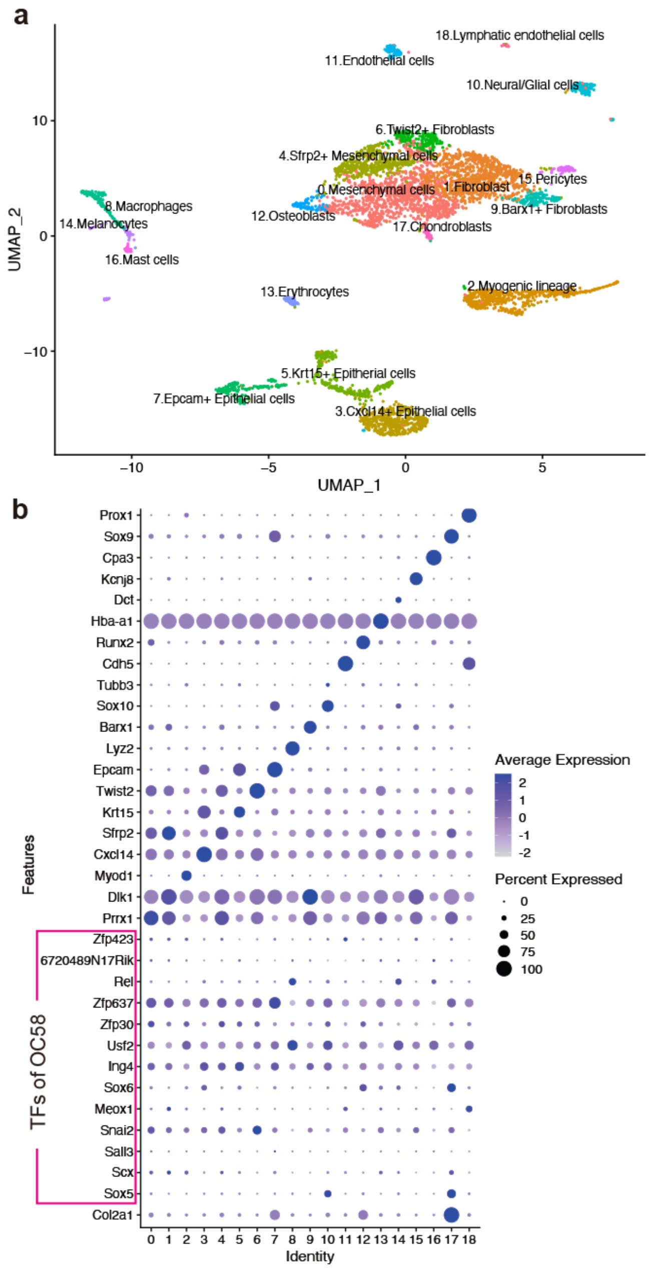


**Supplementary Figure. 17 |** Expression pattern of OC58 in craniofacial cartilage at a single-cell level.

(a) UMAP plot of mouse craniofacial scRNA-seq data at E14.5. Data were from The FaceBase data repository (Record ID 1-DTK2, doi: 10.25550/1-DTK2; http://facebase.org).

(b) Dot plot of marker genes and TFs of OC58. Marker genes are as follows; *Prrx1, Sfrp2* (mesenchymal cells); *Dlk1*(fibroblasts); *Myod1*(myogenic cells); *Cxcl14*(skin epithelium); *Krt15*(epithelial cells); *Twist2*(fibroblasts); *Epcam*(epithelial cells); *Lyz2*(macrophages); *Barx1*(fibroblasts); *Sox10*(neural and glial cells); *Tubb3*(neurons);

*Cdh5*(endothelial cells); *Runx2*(osteoblasts); *Hba-a1*(erythrocytes); *Dct*(melanocytes); *Kcnj8*(pericytes); *Cpa3*(mast cells); *Sox9*(chondroblasts); *Prox1(*lymphatic endothelial cells).
